# End-to-end high-throughput single-cell proteomics via SPRINT and dual-spray LC-MS

**DOI:** 10.1101/2025.11.03.686420

**Authors:** Zhen Liu, Wenbo Dong, Lei Gu, Ruicheng Ge, Xinlei Zeng, Jingwen Deng, Haoran Zhang, Zilu Ye

## Abstract

Single-cell proteomics (SCP) enables direct measurement of protein heterogeneity but remains constrained by throughput and limited applicability to primary tissues. Here, we present an integrated workflow developed to address both challenges. We engineered SPRINT, an AI-powered bioprinting platform that prepares more than 10,000 single cells per day, over tenfold faster than existing systems, while maintaining stability and enabling identification of over 6,000 proteins from individual HeLa cells. To expand analytical capacity, we further designed a dual-spray tandem direct injection (TDI) LC system that parallelizes non-analytical steps with peptide separation, doubling MS utilization efficiency and enabling 168 label-free SCP runs per day without sensitivity loss. Together with optimized tissue-dissociation protocols, this integrated workflow enabled creation of the first mouse tissue-derived SCP atlas, profiling >1,000 single cells across six organs. These advances establish SCP as a scalable platform ready for broad biological and biomedical applications.

## Main

Single-cell proteomics (SCP) enables direct measurement of proteins—the primary effectors of cellular processes—at single-cell resolution. Unlike transcript-based approaches that rely on mRNA as indirect surrogates^1-3^, SCP captures protein-level variation that more faithfully reflects cellular physiology^4-9^, making it a powerful tool for dissecting heterogeneity in complex systems. Recent advances in high-sensitivity workflows have transformed SCP from proof-of-concept to robust applications^6-8^. Miniaturized sample preparation strategies, such as OAD^10^, nanoPOTS^11^, SCoPE-MS^12, 13^, PiSPA^14^, and Chip-Tip^15^, have reduced sample loss and improved peptide recovery, while precision isolation platforms like cellenONE with proteoCHIP further enhanced reproducibility^15, 16^. Combined with next-generation mass spectrometers and DIA-based acquisition strategies^17, 18^, these developments now enable routine detection of thousands of proteins per cell.

Despite these advances, SCP adoption remains limited by unresolved bottlenecks, with throughput as the major constraint. In sample preparation, current workflows typically process only a few thousand cells over several days^19, 20^, during which declining cell viability and variable recovery introduce batch effects and skew population distributions. This low efficiency restricts scalability and makes large studies difficult to execute. At the LC–MS stage, throughput is further constrained by long non-analytical operations— sample loading, column washing, and equilibration—that dominate runtime, leaving most platforms limited to ∼50 single-cell runs per day^15, 21^. Multiplexing strategy^12, 22, 23^ can partially increase throughput but often compromise depth or quantitative accuracy^24, 25^. Together, these constraints create a tradeoff between sensitivity and scale, slowing the transition of SCP from specialized demonstrations to routine biomedical applications. Beyond throughput, the lack of standardized protocols for tissue-derived single cells further hinders progress: most workflows^14, 15, 21^, have been optimized for cultured lines, while primary tissues require tailored dissociation and handling methods to ensure reproducibility and data quality across laboratories.

To overcome these challenges, we developed the Single-cell Proteomics Rapid INTegrator (SPRINT), a dedicated platform designed to dramatically expand the throughput of SCP sample preparation. SPRINT integrates automated, AI-guided dispensing and handling, enabling stable and reproducible processing of more than 10,000 single cells per day. At the analytical stage, we further engineered a custom dual-column tandem direct injection (TDI) system that eliminates non-analytical downtime, effectively doubling MS utilization efficiency and supporting up to 168 label-free single-cell runs daily without compromising sensitivity. In addition, we established optimized tissue-dissociation protocols tailored for SCP, ensuring reproducible proteomic measurements directly from primary cells. Leveraging this integrated workflow—combining SPRINT, the dual-spray TDI system, and tissue-optimized methods—we constructed the first comprehensive tissue-derived SCP atlas, resolving thousands of proteins per cell and uncovering tissue-resident heterogeneity with unprecedented depth. Together, these innovations mark a critical advance, transitioning SCP from proof-of-concept studies to a scalable and broadly applicable platform for biomedical research.

## Results

### SPRINT achieves >10,000 single-cell preparations per day with exceptional stability and sensitivity

Throughput has long been the central barrier to scaling SCP. Current state-of-the-art platforms process only ∼3,000 cells in 1–2 days^19, 20^, limiting studies to small sample sizes and making the analysis of tens of thousands of cells—routine in single-cell transcriptomics—impractical. Prolonged sorting times also risk perturbing cellular states and introducing batch effects. To overcome these constraints, we developed SPRINT, a high-throughput, AI-powered single-cell printing platform that delivers a step-change in SCP sample preparation capacity **(Fig. 1a)**. Built on a multi-nozzle chip with hundreds of independently controlled nozzles, SPRINT can dispense droplets as small as 30 pL, enabling ultra-miniaturized reactions that minimize buffer interference during lysis and digestion. Integrated temperature (4–60 °C) and humidity control (up to 70%) prevent evaporation of trace volumes, while automatic static removal ensures droplets can be precisely deposited at the bottom of well plate. High-resolution imaging combined with AI-based algorithms reliably distinguishes debris, single cells, and doublets, guaranteeing accurate single-cell isolation (**Fig. 1b**).

**Figure 1.**
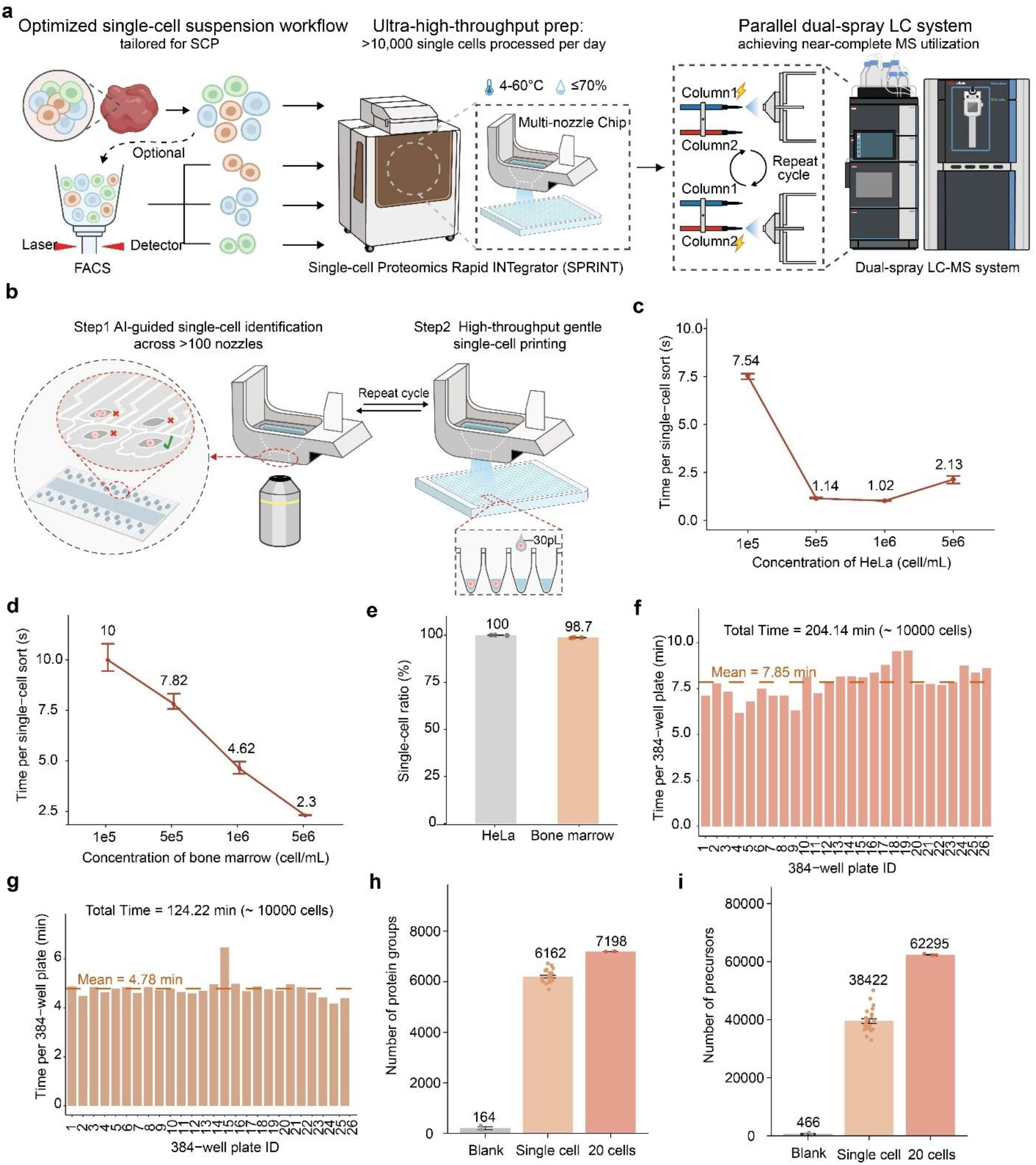
SPRINT achieves >10,000 single-cell preparations per day with exceptional stability and sensitivity. (**a**) Overview of the optimized SCP workflow, integrating high-throughput single-cell preparation (>10,000 cells/day) on the SPRINT platform with a dual-spray LC-MS system for near-complete MS utilization. (**b**) AI-assisted single-cell identification across >100 nozzles (Step 1) and gentle droplet-based single-cell printing (Step 2) using 30 pL droplets. (**c-d**) Sorting time per cell at varying concentrations for HeLa (**c**) and bone marrow (**d**) suspensions. (**e**) Single-cell sorting accuracy for HeLa and bone marrow cells. (**f-g**) Plate-by-plate processing time for ∼10,000 HeLa (**f**) and bone marrow (**g**) cells; mean times per 384-well plate indicated. (**h-i**) Proteome depth from blank controls, single HeLa cells, and 20-cell pools: number of protein groups (**h**) and precursors (**i**) identified by LC-MS.

This combination of innovations allows SPRINT to prepare >10,000 high-quality single-cell samples per day—over tenfold faster than existing systems—while maintaining exceptional robustness across sample types. For HeLa cells, sorting efficiency increased 6.6-fold when cell concentration rose from 1×10^5^ to 5×10^5^ cells/mL, peaking between 5×10^5^ and 1×10^6^ cells/mL (**Fig. 1c**). Bone marrow cells exhibited a linear improvement in efficiency up to 5×10^6^ cells/mL (**Fig. 1d**). Continuous processing of 10,000 cells at optimal concentrations maintained high accuracy—100% for HeLa and 98.7% for bone marrow (**Fig. 1e, Fig. S1**)— at average speeds of 7.85 and 4.78 min per 384-well plate, respectively (**Fig. 1f, g**). Coupled with Vanquish Neo–Orbitrap Astral Zoom LC-MS, SPRINT yielded deep proteome coverage, identifying a median of 6,162 protein groups and 38,422 precursors from single HeLa cells, and over 7,198 protein groups with >62,000 precursors from 20-cell pools (**Fig. 1h, i**). Collectively, these results establish SPRINT as a novel platform that removes the throughput bottleneck in SCP, achieving speed and stability approaching the scale of single-cell transcriptomics while preserving the sensitivity needed for deep proteome profiling.

### Versatile sorting of cultured and tissue-resident cells with SPRINT

Despite the SPRINT platform enabling high-throughput single-cell sorting for cultured and easily obtained tissue cells like bone marrow, isolating other tissue-resident cell populations remains more challenging. This is primarily due to the production of cell debris and the delicate nature of cells during mechanical and enzymatic dissociation, reducing recovery rates and complicating accurate sorting. To address these issues, we enhanced SPRINT sorting algorithm using a shallow neural network for improved debris discrimination (**Fig. 2a**) and developed two interchangeable printing chips tailored to different input conditions (**Fig. 2b**). The regular chip, with 640 independent nozzles, enables rapid sorting at high cell densities, whereas the high-recovery chip features ∼100 nozzles, a shortened loading chamber, and sealed non-working nozzles to concentrate cell flow into the imaging area, improving cell utilization for low-density suspensions. To evaluate printing stability, we systematically analyzed droplet volume consistency by generating 65 million droplets while monitoring two representative nozzles at every 5-million-droplet interval. Both nozzles demonstrated exceptional stability, with median coefficients of variation (CV) of approximately 2% throughout the entire printing process **(Fig. 2c, d)**. Building on this stable performance, we further evaluated the cell-sorting efficiency by comparing the two chip designs. The high-recovery chip improved sorting efficiency for low-density samples, outperforming the regular chip when processing 50 μL of 1×10^5^ cells/mL HeLa suspensions (**Fig. 2e**). We next assessed cell recovery at low input numbers and sorting efficiency across different cell concentrations using tissue-derived lymph, brain, and intestinal cell suspensions. In recovery tests with 500 input cells (5 μL at 1×10^5^ cells/mL), the high-recovery chip consistently yielded higher recovery rates than the regular chip **(Fig. 2f-h)**. Moreover, sorting efficiency increased with cell concentration across all these cell types **(Fig. 2i-k)**. These results demonstrate SPRINT’s versatility in efficiently sorting both cultured and diverse tissue-derived cell types, from high-yield preparations to scarce, low-density suspensions.

**Figure 2.**
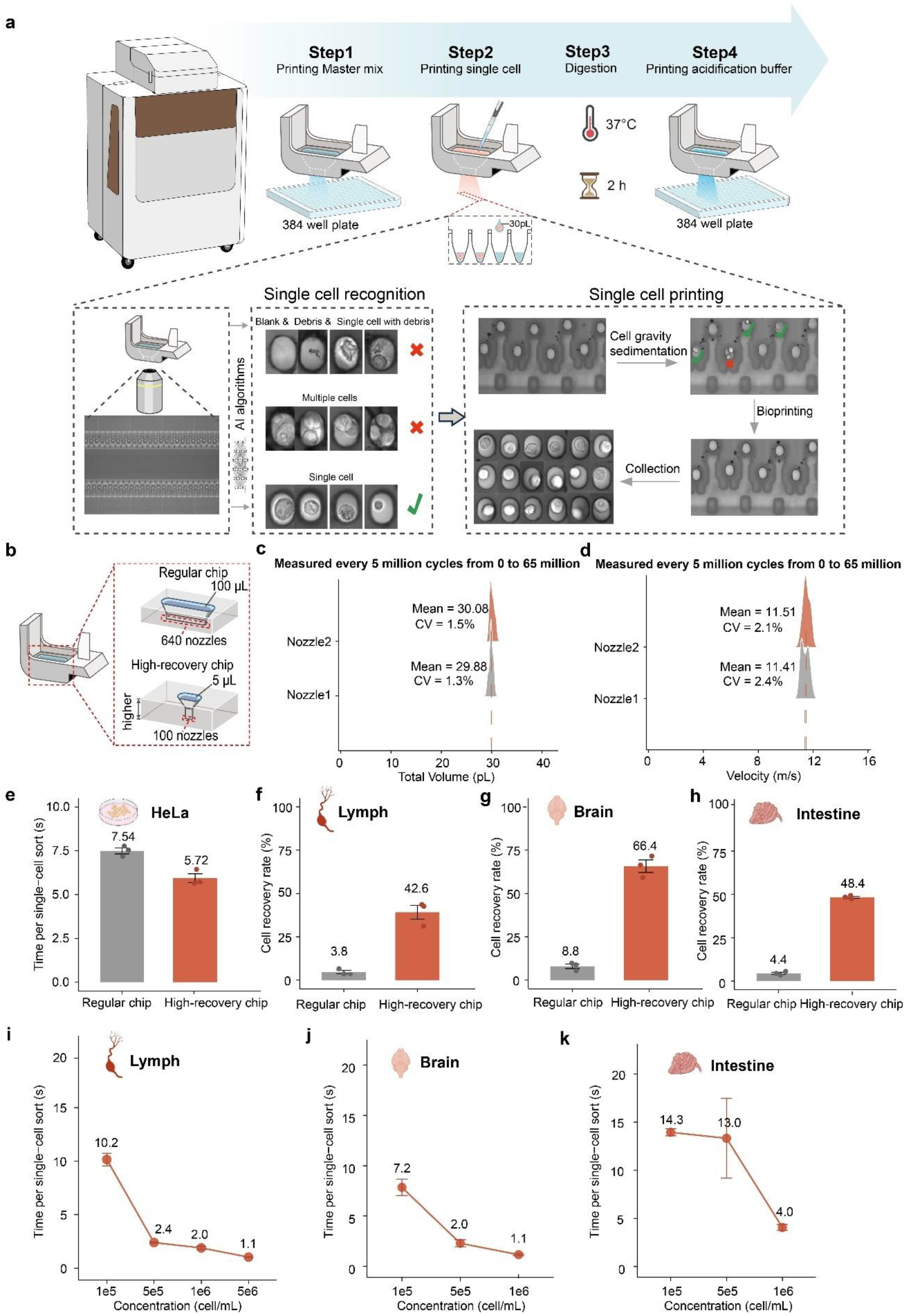
Versatile sorting of cultured and tissue-resident cells with SPRINT. (**a**) Detailed workflow for single-cell sample preparation using the SPRINT platform, incorporating AI-based single-cell recognition and gentle droplet printing. (**b**) Design of the regular chip (640 nozzles, 100 µL loading) and high-recovery chip (∼100 nozzles, 5 µL loading) optimized for low-density samples. (**c-d**) Droplet volume (**c**) and velocity (**d**) stability over 65 million dispensing cycles for two representative nozzles. (**e**) Sorting time per HeLa cell for regular and high-recovery chip at 50 µL of 1×10^5^ cells/mL. (**f-h**) Recovery rates for 500 cells from lymph (**f**), brain (**g**), and intestine (**h**) suspensions using regular vs. high-recovery chip. (**i-k**) Sorting time per cell at increasing concentrations for lymph (**i**), brain (**j**), and intestine (**k**) suspensions using the regular chip.

### Enhanced single-cell proteomics depth through integrated workflow optimization

While the advances in SCP towards an era of deep proteome coverage^14, 15, 21, 26, 27^, SCP workflows still vary considerably across laboratories, lacking standardized protocols. To address this, we leveraged the high-throughput and reproducible SPRINT platform to systematically optimize each step of sample preparation, establishing a robust, standardized workflow for SCP (**Fig. 3a**). We first examined the impact of master mix volume (100-1,000 nL) on proteome coverage by printing individual cells into 384-well plates and analyzing them with a Vanquish Neo–Orbitrap Astral system (**Fig. 3b, Fig. S2a)**. Protein identifications increased sharply between 100 and 300 nL, plateaued between 300 and 500 nL, and declined by 6.9% at 1,000 nL—likely due to protein adsorption losses. We therefore selected 500 nL as the optimal volume. Next, we compared digestion enzymes: 20 ng/μL trypsin, 20 ng/μL Lys-C, or a 1:1 mixture (10 ng/μL each). Trypsin alone yielded the highest protein and precursor identifications (**Fig. 3c, Fig. S2b**), whereas Lys-C alone produced longer peptides less suited for MS detection. Trypsin was thus established as the optimal protease. We next investigated the impact of digestion time (0.5-4 h) using 500 nL of 20 ng/μL trypsin. This analysis revealed pronounced time-dependent variations in proteome coverage (**Fig. 3d, Fig. S2c**). While 0.5 h digestion achieved comparable protein identification to 1-2 h treatments, it exhibited significantly higher missed cleavage rates (**Fig. 3e**), indicating incomplete digestion. Notably, extended 4 h digestion showed a paradoxical 19.7% reduction in protein identifications despite achieving optimal cleavage efficiency (**Fig. 3e**), likely due to evaporation. An incubation of 1–2 h provided the best balance of completeness and coverage. We further assessed acidification and ionization conditions. Comparing acid types, 0.1% trifluoroacetic acid (TFA) in 1% DMSO improved protein group identifications by 9.2% (single-cell) and 14.3% (20-cell) relative to 0.1% formic acid (FA) (**Fig. 3f**), and mitigated precursor loss seen with FA (**Fig. S2d**). Low concentrations of DMSO can enhance peptide ionization^28, 29^, but the optimal level for SCP was unclear. To systematically evaluate its effect on SCP sensitivity, we tested DMSO concentrations ranging from 0% to 5% in 0.1% TFA buffer (**Fig. 3g, Fig. S2e**). Increasing DMSO from 0% to 0.5% in 0.1% TFA buffer improved identifications, whereas 1% DMSO slightly reduced them, especially for precursor counts in 20-cells samples (**Fig. 3g, Fig. S2e**). We therefore set 0.5% DMSO as optimal^15, 26^. Finally, we evaluated levels of carrier proteome which was used for spectral library generation in label-free SCP^15^, sorting 10-100 cells per well. Identification depth was similar across all carrier amounts (**Fig. 3h, Fig. S2f**), indicating that 10-20 carrier cells are sufficient for conventional SCP without compromising quality. Together, these optimizations define an integrated SCP sample preparation workflow that maximizes proteome depth while maintaining high reproducibility, providing a standardized foundation for cross-study comparability.

**Figure 3.**
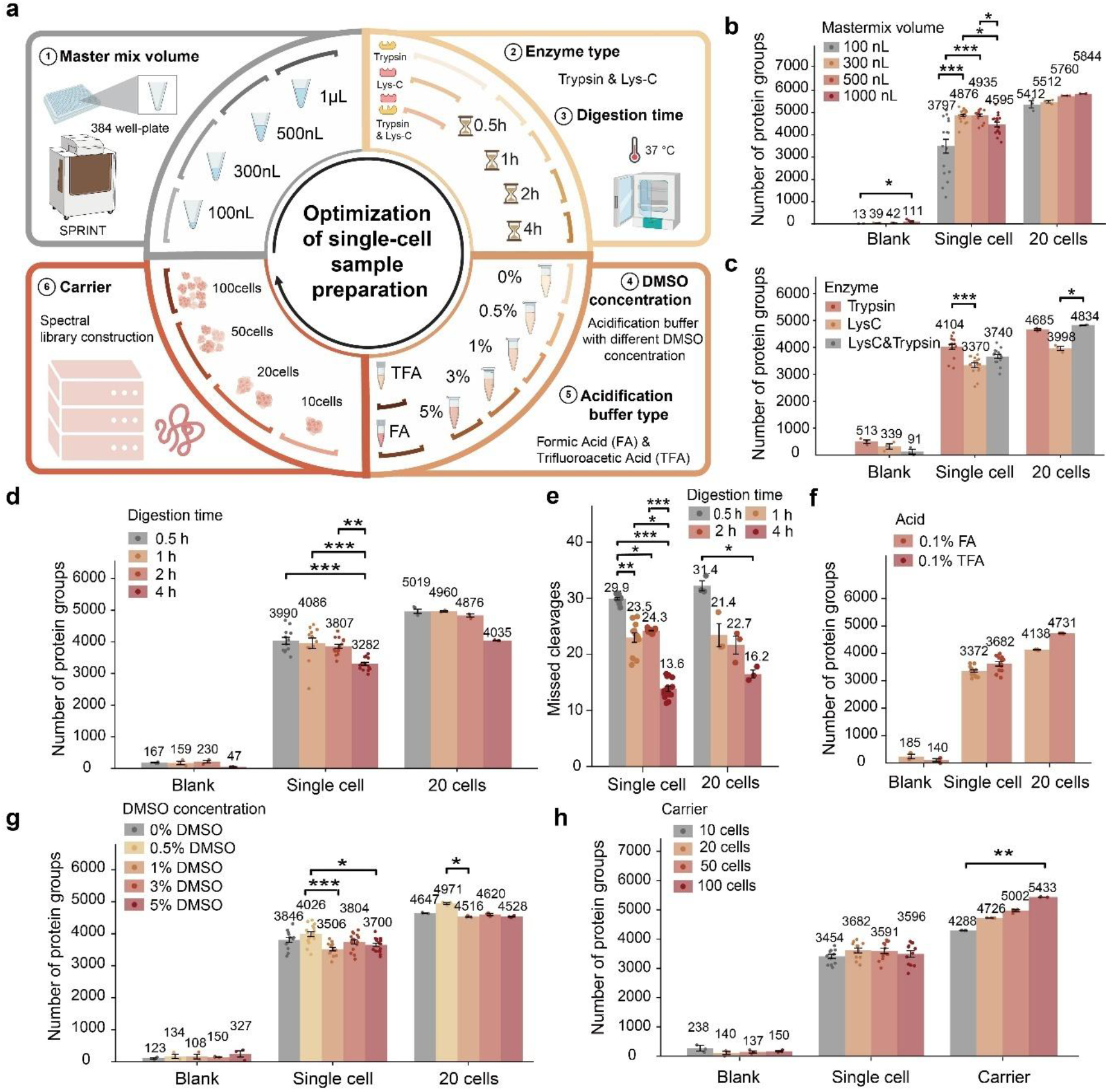
Enhanced single-cell proteomics depth through integrated workflow optimization. (**a**) Schematic of the optimization framework, including master mix volume, enzyme type, digestion time, acidification buffer, DMSO concentration, and carrier cell load. (**b**) Protein identifications across different master mix volumes. (**c**) Effect of enzyme type on protein identifications. (**d**) Impact of digestion time on protein identifications and (**e**) corresponding missed cleavage rates. (**f**) Comparison of 0.1% FA versus 0.1% TFA in 1% DMSO buffer. (**g**) Effect of DMSO concentration on protein identifications. (**h**) Effect of carrier cell number on protein identifications. Data represents HeLa single-cell or 20-cell analyses, with blanks as controls. *, p < 0.05; **, p < 0.01; ***, p < 0.001. Except for panel f (Wilcoxon rank-sum test), statistical analyses used Dunn’s test following Kruskal-Wallis analysis with Bonferroni-adjusted p-values.

### Comprehensive tissue dissociation strategy tailored for single-cell proteomic analysis

With the optimized SPRINT workflow enables deep and consistent proteome coverage for cultured cells, we next aimed to develop a robust and efficient protocol for tissue-derived single cells. Existing single-cell transcriptomics dissociation methods^30-32^ are insufficient for proteomics, as they prioritize RNA preservation without addressing protein-specific issues such as cellular debris interference. To address this gap, we established a tissue dissociation protocol specifically optimized for SCP (**Fig. 4a**). After anesthesia, we collected 12 mouse biospecimens including brain, intestine, lymph, spleen, thymus, bone marrow, heart, kidney, liver, lung, testis, and peripheral blood and prepared single-cell suspensions using our optimized workflow (**Fig. 4b-m**) with systematic evaluation of bovine serum albumin (BSA), a common additive in transcriptomics^30, 33^, for its role in SCP.

**Figure 4.**
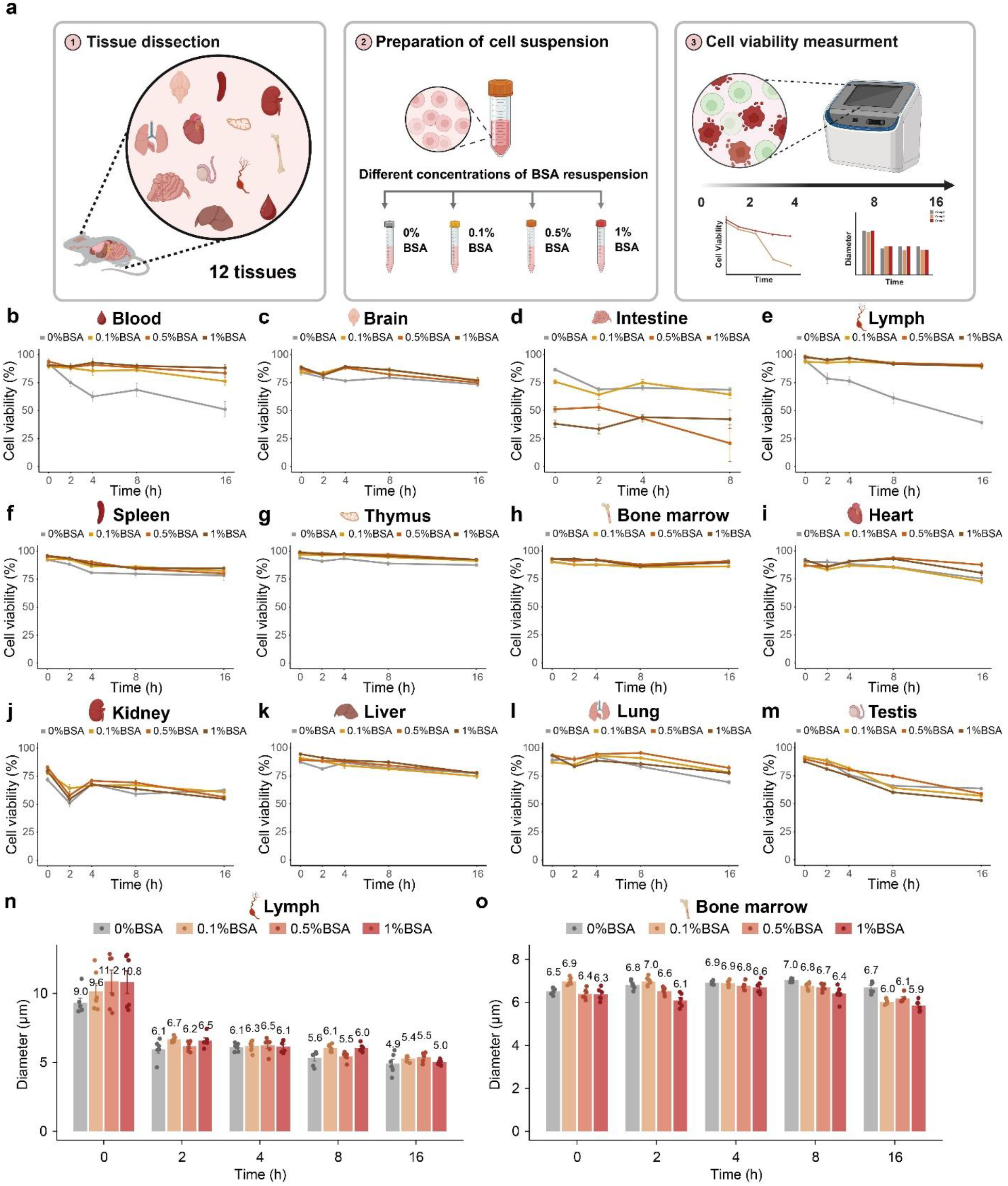
Comprehensive tissue dissociation strategy tailored for single-cell proteomic analysis. (**a**) Workflow for optimizing tissue-derived single-cell preparation, including tissue dissection, suspension in buffers with varying BSA concentrations, and cell viability assessment. (**b-m**) Time-course viability analysis of single cells from blood (**b**), brain (**c**), intestine (**d**), lymph (**e**), spleen (**f**), thymus (**g**), bone marrow (**h**), heart (**i**), kidney (**j**), liver (**k**), lung (**l**), and testis (**m**) tissues under different BSA concentrations. (**n-o**) Cell diameter changes over time for lymph (**n**) and bone marrow (**o**) cells across tested conditions.

We conducted a time-course assessment by partitioning cell suspensions into aliquots with varying BSA concentrations (0-1%), except brain cells, which were suspended in HBSS. Viability was measured at 0, 2, 4, 8, and 16 h. Lymph and blood cells showed strong protection, maintaining ∼90% and ∼80% survival at 16 h versus 40-50% in PBS (**Fig. 4b, e**). Brain, spleen, and thymus cells exhibited moderate protection (**Fig. 4c, f, g**). In contrast, intestine cells were highly sensitive: BSA reduced viability in a concentration-dependent manner, with near-complete death by 8 h at ≥0.5% (**Fig. 4d**). BSA also induced morphology changes (**Fig. 4n-o, Fig. S3**). Lymph and thymus cells first swelled then shrank (**Fig. 4n, Fig. S3b**), intestine cells shrank with higher BSA (**Fig. S3e**), and liver cells enlarged over time (**Fig. S3g**). Other tissues showed modest or mixed effects (**Fig. S3**). Next, we conducted an investigation to determine whether BSA affects SCP. HeLa cells were suspended in varying BSA concentrations and then subjected to Vanquish Neo– Orbitrap Astral system analysis. We found that 1% BSA produced high background and significantly reduced protein identifications, while 0.1-0.5% gave comparable results (**Fig. S4-S5**). We therefore standardized on 0.1% BSA for SCP workflows. By systematically evaluating viability, morphology, and proteomic compatibility, we establish practical guidelines for tissue dissociation in SCP. These results emphasize that dissociation buffers must be tailored to tissue type and that low BSA concentrations provide a balance between stabilizing cells and ensuring reliable proteomic analysis.

### Maximizing MS utilization efficiency via a dual-spray TDI system

The SPRINT platform enables preparation of tens of thousands of SCP samples within hours. However, sample throughput now exceeds instrument capacity^15, 21^, as the main bottleneck lies not in MS acquisition speed but in the LC step. A full LC cycle comprised loading, washing, equilibration and analyte migration, with most of these steps representing LC system preparation during which the mass spectrometer remains idle. To overcome this, we developed a dual-spray TDI system that operates two independent analytical columns in parallel, allowing one to equilibrate and load while the other performs peptide separation (**Fig. 5a**). A custom-built controller manages valve switching, column positioning, voltage, and temperature, maintaining compatibility with the Vanquish Neo LC and Orbitrap Astral MS. We define MS utilization efficiency as the fraction of total runtime during which the MS actively acquires peptide data, reflecting both LC and MS performance. Using 100 ng HeLa digests, both columns delivered highly consistent performance, enabling the identification of ∼6,880 proteins and ∼115,000 precursors each (**Fig. 5b, c**), with excellent chromatographic reproducibility (retention time deviation <0.25 min, **Fig. 5d**), consistent peak areas (**Fig. 5e**), and robust protein quantification confirmed by principal component analysis (PCA) (**Fig. 5f**). Additional benchmarks demonstrated minimal batch effects and high reproducibility across replicate runs (**Fig. S6**).

**Figure 5.**
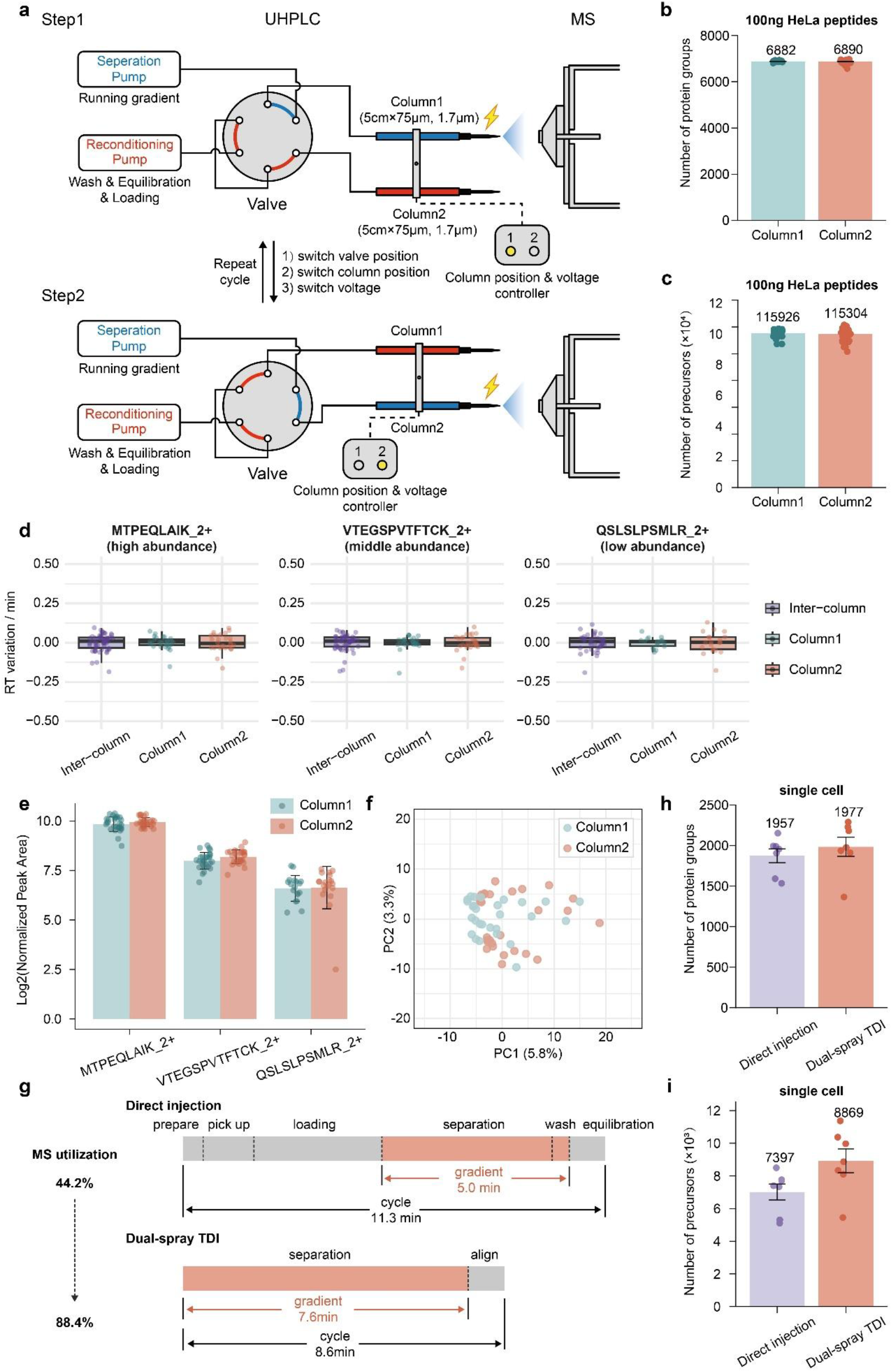
Maximizing MS utilization efficiency via a dual-spray TDI system. (**a**) Schematic of the dual-spray TDI workflow, featuring tandem LC, two 5 cm× 75 μm columns, and a custom-built column controller for parallel loading and separation. Proteins (**b**) and precursors (**c**) identified from 100 ng HeLa peptides analyzed on both columns using a 12-min cycle method. (**d-e**) Retention time variation (**d**) and normalized peak areas (**e**) of representative precursors from high-, medium-, and low-abundance ranges across columns. (**f**) PCA showing reproducible protein quantification between columns. (**g**) Comparison of MS utilization efficiency: conventional direct injection (11.3-min cycle, 44.2%) versus dual-spray TDI (8.6-min cycle, 88.4%). (**h-i**) Single-cell analysis of embryonic day 10.5 mouse cells shows comparable sensitivity between configurations, with proteins -(**h**) and precursors (**i**) consistently identified.

After confirming the system’s robustness with bulk samples, we proceeded to test its capabilities in SCP. Using our optimized 8.6-minute gradient, 7.6 minutes are dedicated to effective peptide separation, resulting in an MS utilization efficiency of 88.4% (**Fig. 5g**). In comparison, the conventional single-spray configuration requires 11.3 minutes to complete a 5-minute separation, with over half of the cycle time consumed by sample loading, washing, equilibration and analyte migration—yielding a utilization efficiency of just 44.2% (**Fig. 5g**). This two-fold improvement in MS utilization efficiency highlights the superior performance of system for short-gradient analysis, enabling sensitive proteomic analysis at a throughput of proximately 168 label-free SCP samples per day. We further validated the applicability of the dual-spray TDI system in SCP using embryonic day (E) 10.5 mouse cells as a model. Comparative analysis demonstrated that the dual-spray configuration achieved higher identifications than the conventional single-spray, with median protein counts of 1,977 and 1,957, respectively, and precursor counts of 8,869 and 7,397, respectively (**Fig. 5h, i**). Together, we report the first dual-spray TDI system tailored for SCP, overcoming LC bottlenecks without compromising sensitivity. By parallelizing column loading, washing, equilibration and analyte migration with separation, the system nearly doubles throughput and achieves up to 88.4% MS utilization efficiency, with potential to approach 100%. This establishes a robust and scalable path toward truly high-throughput SCP.

### Single-cell proteomics atlas of mouse reveals novel dimensions of cellular heterogeneity

Benefiting from the overall workflow optimization described above, we constructed a high-quality SCP atlas across diverse mouse tissues, including bone marrow, heart, intestine, kidney, lung, and lymph (**Fig. 6a**). Mouse tissues were dissociated into single-cell suspensions using optimized protocols and sorted with SPRINT, which also provided AI-based cell diameter measurements. Bone marrow and lymph showed lower CVs, consistent with their immune-cell homogeneity, while intestine displayed higher CVs due to the fragility of its cells (**Fig. S7**). High-quality single-cells were then analyzed by high-throughput dual-spray TDI system, leading to the identification of 5,823 unique proteins (**Fig. 6b**). For cells from different tissues, the median identification ranged from 559 to 1,052 protein groups for single-cells, and from 1,722 to 2,663 protein groups for 20-cells pool (**Fig. S8**). Of the 5,823 identified proteins, 1,039 were commonly identified across all six tissues (**Fig. 6b**). These shared proteins largely comprise core cellular components, including ribosomal proteins, metabolic enzymes, and cytoskeletal elements. Conversely, tissue-specific proteins, such as cardiac troponins and pulmonary surfactant-associated proteins, exhibited restricted expression in their corresponding tissues^34^.

**Figure 6.**
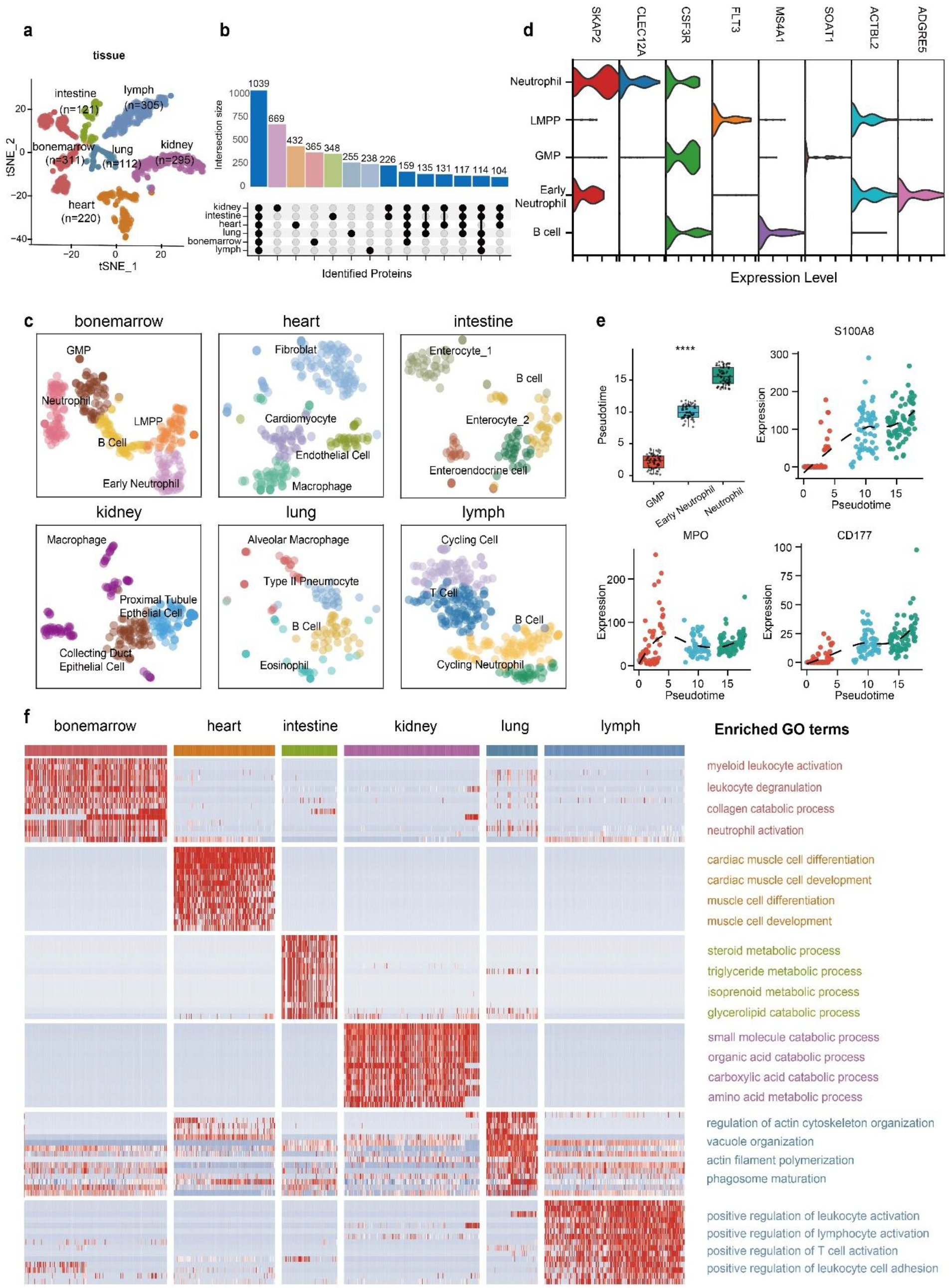
Single-cell proteomics atlas of the mouse reveals novel dimensions of cellular heterogeneity. (**a**) t-SNE visualization of single-cell proteomes from bone marrow, heart, intestine, kidney, lung, and lymph, with cells colored by tissue origin. (**b**) UpSet plot showing overlap of proteins identified across tissues at the single-cell level. (**c**) Unsupervised clustering with marker-based annotation reveals major tissue-resident populations. (**d**) Violin plots of canonical marker-protein expression confirming cell-type identities. (**e**) Pseudotime trajectory analysis of neutrophil differentiation from GMPs to mature neutrophils, showing stage-specific marker expression. (**f**) Heatmap of tissue-enriched marker proteins and their associated GO biological processes.

To resolve cellular heterogeneity, unsupervised clustering and marker-based annotation identified major tissue-resident populations, including cardiomyocytes (heart) ^38^, enterocytes and enteroendocrine cells (intestine) ^39^, proximal tubule and collecting duct epithelial cells (kidney) ^40^, and eosinophils and type II pneumocytes (lung) ^41^ (**Fig. 6c**). Canonical marker-protein expression patterns clearly confirmed the identity of these cell populations (**Fig. 6d, Fig. S9**). Single-cell analysis revealed tissue-specific functional associations, such as metabolic processes in kidney epithelial cells and immune responses in lymph B cells, alongside roles in tissue organization (**Fig. S10-S11**). An interesting example comes from neutrophils, the most abundant leukocytes^42, 43^. Pseudotime analysis traced their progression from granulocyte–monocyte progenitors to maturity (**Fig. 6e**). SCP resolved intermediate states and stage-specific protein expression, with S100A8 and CD177 rising during maturation^44, 45^ and MPO peaking early. Unlike bulk or scRNA-seq^42^, SCP captured dynamic protein-level variations, underscoring its power in dissecting neutrophil differentiation. Functional enrichment analysis of the most highly expressed proteins in each tissue revealed pathways consistent with the known biological functions of the corresponding organs (**Fig. 6f**). For example, top expressed proteins in the heart were enriched for cardiac muscle development^35^, while those in the intestine were enriched for lipid and isoprenoid metabolic processes^36^. Similarly, proteins highly expressed in bone marrow and lung were associated with immune cell functions, showing enrichment in pathways such as neutrophil degranulation, leukocyte activation, and phagosome maturation^37, 38^. This SCP atlas provides both a valuable resource and a methodological guideline for dissecting tissue organization, cellular heterogeneity, and differentiation dynamics at the protein level across multiple mouse organs.

## Discussion

SCP has entered a stage where sensitivity no longer represents the primary limitation. With optimized sample processing strategies and next-generation mass spectrometers, the routine identification of thousands of proteins per single cell has become possible. What continues to hinder broad adoption is scalability. Throughput constraints in both sample preparation and LC–MS analysis, together with the lack of standardized protocols for tissue-derived cells, have prevented SCP from being widely applied to diverse biological and clinical questions.

In this study, we addressed these bottlenecks through an integrated and end-to-end workflow. Central to this advance is SPRINT, an AI-powered printing platform that prepares more than 10,000 single-cell samples per day. This represents a >tenfold increase in throughput compared with existing SCP systems, which typically handle on the order of ∼1,000 cells per day. By dramatically shortening preparation time, SPRINT also minimizes cell viability loss during extended handling, enabling robust analysis of fragile tissue-derived cells. Importantly, this level of throughput brings SCP much closer to the scale of single-cell transcriptomics, positioning it as the first proteomics platform approaching the efficiency of scRNA-seq workflows. Next, we introduced a TDI LC system, the first of its kind for SCP. Previous multicolumn or dual-spray approaches have been explored to increase throughput, but they often rely on trap-elution configurations, require long columns, and deliver only modest gains in effective MS usage, making them poorly suited for SCP^39-44^. In contrast, our system parallelizes loading, washing, equilibration, and analyte migration with peptide separation, doubling MS utilization efficiency and enabling 168 label-free single-cell analyses per day without compromising sensitivity. Leveraging these technical innovations, we generated a tissue-derived single-cell proteome atlas in mouse, covering six organs and over 5,800 proteins. This atlas resolved major tissue-resident cell populations, captured functional pathways aligned with organ physiology, and dissected differentiation trajectories such as neutrophil maturation with unprecedented resolution. Beyond serving as a resource, these findings demonstrate the readiness of SCP to address fundamental biological and biomedical questions.

Looking forward, we envision that further improvements in ion source design and LC configuration will be essential to push daily throughput for label-free SCP into the range of 300–500 cells, enabling studies at a scale relevant for population-level biology. In parallel, the development of new labeling chemistries and multiplexing reagents may drive even more sensitive and scalable SCP, ultimately narrowing the gap with single-cell transcriptomics in terms of throughput. Beyond hardware and chemistry, advances in automated data analysis pipelines, real-time quality control, and multi-omics integration will be critical to ensure reproducibility and accessibility across laboratories. At the same time, the establishment of clinical-grade dissociation and tissue-handling protocols will lay the foundation for translating SCP into biomedical and therapeutic contexts. Together, these directions point toward the broader vision of building a virtual cell—a comprehensive molecular model that integrates proteomic, transcriptomic, and spatial dimensions to capture cell states and dynamics at unprecedented resolution.

In summary, by combining SPRINT, the dual-spray TDI system, and tissue-optimized protocols, we present an integrated SCP workflow that simultaneously achieves high sensitivity, high throughput, and applicability to primary tissues. This work represents a decisive step in establishing SCP as a scalable and broadly applicable platform for the life sciences.

## Material and methods Cell culture

The HeLa cells were grown in DMEM medium containing 10% fetal bovine serum, 1% penicillin and streptomycin at 37 °C in a humidified atmosphere with 5% CO_2._ The cells were washed three times with PBS. Cells were then resuspended in PBS at 1000 cells/µL with varying BSA concentration for cell sorting within the SPRINT.

### Mice and Tissue dissociation

The mouse work was performed in accordance with the Institutional Animal Care and Use Committee (IACUC) of Suzhou Institute of Systems Medicine (ISM-IACUC-0151-R), and animals were housed in specific-pathogen-free mouse facilities. The 8-week-old male C57BL6/J mice were housed in standard conditions, with 12-h/12-h light/dark cycles and a controlled temperature of 22-24 °C and humidity of 60%, with unrestricted food and water availability, and were examined daily.

All tissue and organ collections were performed in full compliance with institutional ethical standards for animal experimentation. Mice were anesthetized with 2.5% tribromoethanol administered intraperitoneally at a dosage of 10 μL per gram of body weight. Following anesthesia, cardiac perfusion was carried out using ice-cold PBS to ensure thorough blood removal prior to tissue harvesting. Twelve tissues and organs were collected from mice for subsequent cell dissociation, including the heart, liver, lung, brain, intestine, lymph, thymus, spleen, kidney, testes, bone marrow, and blood. Due to the high amount of cellular debris generated during the dissociation of brain, liver, and kidney tissues, an additional debris removal step using Debris Removal Solution (Miltenyi Biotec) was performed to ensure cleaner single-cell suspensions. The specific dissociation procedures for each tissue are described as follows:

Solid tissues, with the exception of the brain, were rinsed 3-5 times with PBS containing 0.04% BSA. Brain tissue was rinsed 2-3 times with dulbecco’s phosphate-buffered saline (DPBS) supplemented with 0.04% BSA. After washing, an appropriate volume of RPMI 1640 medium (Thermo Fisher Scientific) with 0.04% BSA was added, and the tissues were finely minced into approximately 0.5 cm^3^ pieces using sterilized scissors. The minced tissues were then subjected to enzymatic digestion for subsequent single-cell preparation.

Unlike solid tissues, bone marrow and blood required distinct processing methods. Bone marrow was flushed out from the cleaned femurs using a syringe filled with RPMI 1640 supplemented with 0.04% BSA. Freshly collected blood was obtained via ocular puncture and transferred into EDTA·K2-coated anticoagulant tubes, followed by gentle mixing. Bone marrow and blood samples were first passed through a 70 μm cell strainer. Red blood cells were then lysed using Red Cell Lysis Buffer at a 1:3 (sample: buffer) ratio, with bone marrow samples incubated for 5 min and blood samples for 10 min at room temperature. The lysis reaction was terminated by adding an equal volume of PBS, and cells were pelleted by centrifugation at 500 g for 5 min at 4 °C. The resulting cell pellets were washed using RPMI 1640 with 0.04% BSA prior to further use.

For single-cell preparation from the brain, papain and DNase I (both from Worthington) were first dissolved in Hibernate A medium (Gibco) and incubated at 37 °C on a rotator (10 rpm) for 15 min to generate the enzyme stock solution. This stock was then cooled on ice and diluted in RPMI 1640 supplemented with 0.04% BSA to final concentrations of 1 μg/μL for papain and 0.1 μg/μL for DNase I, yielding the digestion solution. The minced brain tissue was then added to the digestion solution and incubated at 37 °C with gentle agitation on a thermostatic shaker at 500 g for 15 min. After complete digestion, the cell suspension was passed through a 70 μm cell strainer to remove undigested debris. Cells were subsequently collected by centrifugation at 500 g for 5 min at 4 °C. For debris removal, the cell pellet was resuspended in a pre-chilled PBS and Debris Removal Solution mixture (10:3, v/v; Miltenyi Biotec), following the manufacturer’s instructions. The suspension was carefully overlaid with 4 mL of cold PBS and subjected to density gradient centrifugation at 3,000 g for 10 min at 4 °C with maximum acceleration and deceleration settings. The upper two debris-rich layers were discarded, and the remaining phase was diluted with cold PBS to a final volume of 15 mL. After gentle mixing by inversion, the cells were centrifuged at 500 g for 10 min at 4 °C, and the supernatant was completely removed. After debris removal, 1mL RPMI 1640 medium supplemented with 0.04% BSA was added to the cell suspension and mixed thoroughly. The cells were then washed by centrifugation at 300 g for 5 min at 4 °C to remove residual debris removal solution.

For single-cell preparation from the heart, lung, and testes, collagenase I (Thermo Fisher Scientific) was first dissolved in calcium- and magnesium-free Hanks’ Balanced Salt Solution (HBSS, Gibco) to prepare a stock solution. This was then diluted with RPMI 1640 medium containing 0.04% BSA to a final concentration of 2 μg/μL. DNase I was added directly to the digestion solution at a final concentration of 0.1 μg/μL to prevent cell aggregation. For the heart and liver, digestion was performed at 37 °C on a thermostatic shaker at 500 g for 30 min, whereas for the testes, digestion was limited to 20 min. To further enhance tissue dissociation, 0.25% trypsin was added, followed by an additional incubation at 37 °C, 500 g for 15 min. After enzymatic digestion, the cell suspension was passed through a 70 μm cell strainer and centrifuged at 500 g for 5 min at 4 °C. Red blood cells were removed using the above methods with a 5-min incubation at room temperature, and cell pellets were subsequently washed also using RPMI 1640 medium supplemented with 0.04% BSA to obtain clean single-cell suspensions.

For kidney, spleen, lymph, and thymus, single-cell suspensions were prepared by enzymatic digestion with collagenase II (Thermo Fisher Scientific), pre-dissolved in calcium- and magnesium-free HBSS (Gibco) to generate a stock solution. The working digestion solution was prepared by diluting the stock in RPMI 1640 medium with 0.04% BSA to a final collagenase concentration of 2 μg/μL, followed by the addition of DNase I to 0.1 μg/μL. Tissues were digested at 37 °C and 500 g for 15 min (30 min for kidney), then further treated with 0.25% trypsin under the same conditions for 15 min to enhance dissociation. Cell suspensions were filtered through a 70 μm strainer and centrifuged at 500 g, 4 °C for 5 min. Red blood cells were lysed at room temperature for 5 min as described above. Due to higher debris content, kidney-derived suspensions underwent an additional cleanup step using the same debris removal method as for brain tissue, followed by an extra filtration through a 40 μm strainer and washing with RPMI 1640 containing 0.04% BSA.

For single-cell preparation from liver and intestinal tissues, minced samples were enzymatically digested using a two-step procedure. First, collagenase IV was dissolved in calcium- and magnesium-free Hanks’ Balanced Salt Solution (HBSS, Gibco) to prepare an enzyme stock solution. This stock and DNase I was then diluted in RPMI 1640 medium supplemented with 0.04% BSA to final concentrations of 2 μg/μL (collagenase IV) and 0.1 μg/μL (DNase I) to create the digestion solution. Tissues were digested at 37 °C and 500 g for 20 mins, followed by an additional 15-minute digestion with 0.25% trypsin under the same conditions. Cell suspensions were then passed through a 70 μm strainer and centrifuged at 500 g for 5 min at 4 °C to pellet the cells. Red blood cells were lysed by incubating the samples with Red Cell Lysis Buffer at room temperature for 5 min. To further enhance single-cell purity, liver-derived suspensions underwent an additional debris removal step using a gradient centrifugation method as described for brain tissue.

The procedure for isolating single-cell suspensions from mouse embryos is generally similar to that used for adult mouse tissues, with a key difference in the method of euthanasia. Pregnant mice at embryonic day 10.5 (with the day of vaginal plug detection designated as embryonic day 0.5) were euthanized by cervical dislocation. The uterus was rinsed with sterile saline, and embryos were dissected under a dissection microscope in DPBS buffer. Following collection, the embryos were finely minced and enzymatically dissociated using collagenase I for 30 min and trypsin for 15 min as previously described. The resulting cell suspension was passed through a 70 μm cell strainer to remove debris. Red blood cells were lysed using Red Cell Lysis Buffer, and the remaining cells were washed thoroughly with RPMI 1640 medium supplemented with 0.04% BSA for further use.

### Cell viability and diameter measurement of tissue cells

HeLa cells and different tissue cells were enriched by centrifugation at 500 g for 5 min at 4 °C and then were separately resuspended in PBS buffer containing 0%, 0.1%, 0.5%, and 1% BSA, except for brain cells, which were suspended in HBSS with different BSA concentrations. The cell suspensions with varying BSA concentrations were equally divided into four parts, and cell viability and diameter were measured at 0 h, 2 h, 4 h, 8 h, and 16 h, respectively. Briefly, an equal volume of cell suspension and AO/PI (acridine orange/propidium iodide) solution was mixed and loaded onto a cell counting plate, followed by viability and diameter detection using an automated cell counter.

### Stability analysis of printing chip

The droplet volume and velocity were measured using a Dropwatcher system, which was triggered synchronously with the droplet firing signal. Based on stroboscopic imaging principles, the device captures stable images of ejected droplets within its field of view. Droplet volume is calculated through image analysis of the droplet size. For velocity measurement, the system sequentially captures two droplet images at a fixed time interval (e.g., 15 μs) and computes the displacement between them. The velocity is then derived from this displacement and the known time interval through computational analysis.

### Sample preparation in SPRINT

The SCP sample preparation was performed using a standard 384 well plate (Eppendorf twin.tec® PCR Plate 384 LoBind). Firstly, it needs to calibrate the equipment by using the red ink, where was added to the chip inside the SPRINT. The system was suitable for SCP analysis when the droplet could be printed into the bottom center of the 384 well plate. After calibration, the static electricity was removed to guarantee the printing stability. Then the new prepared 384 well plate was placed inside the SPRINT, where 0.5 µL of master mix buffer (0.2% DDM (D4641-500MG, Sigma Aldrich, Germany), 100mM TEAB, 20 ng/µL trypsin) was automatically printed into each well of the 384 well plate. After that, the chip inside the SPRINT was washed three times with H2O and PBS, respectively. The 1,000 cells/µL was added to the chip, and the single-cell was sorted by high-definition imaging and artificial intelligence algorithms. After sorting, the 384 well plate was then subjected to a controlled incubation phase at 50 °C with 85% relative humidity for 2 h within the instrument environment. The automatic cycle system involved the addition of 500 nL of water to each well until the process was complete. After incubation, the temperature was reduced to 20 °C to stabilize the conditions post-reaction. The chip was washed three times H2O and PBS, respectively. The digestion reaction was stopped by printing 2 µL of 0.1% TFA/0.5% DMSO into the wells. The plates were then sealed with matching 384 well plate covers and placed in the Vanquish Neo, allowing for the direct injection of 2.5 µL ready for MS analysis. For single-cell sample preparation from different tissues, the preparation process was same as the above mentioned.

### Condition optimization for single-cell samples preparation

To systematically optimize SCP workflows, we first evaluated different master mix volumes (100, 300, 500, and 1000 nL) dispensed into 384-well plates using a regular chip, followed by single-cell sorting and digestion termination. Enzyme conditions were tested by dispensing 500 nL of 20 ng/μL trypsin, 20 ng/μL Lys-C, or a 10 ng/μL trypsin/Lys-C mixture. Digestion time was optimized across 0.5, 1, 2, and 4 h after trypsin dispensing (500 nL, 20 ng/μL) and cell sorting.

For acid conditions optimization, digestion was terminated with 1% DMSO containing 0.1%FA or 1%TFA, while DMSO concentration were compared using 0.1% TFA in 0-5% DMSO. Carrier cell effects were assessed by sorting 10-100 cells per well. All reactions were stopped after 2 h (unless testing time courses) with 2 μL of termination solution (0.1% TFA/0.5% DMSO unless otherwise specified), followed by plate sealing and direct 2.5 μL injection into the Vanquish Neo for MS analysis.

### Comparative analysis of cell printing performance between regular chip and high recovery chip

Cell sorting efficiency was evaluated using HeLa cells, lymph cells, brain cells and intestine cells. HeLa cell suspensions were prepared at concentrations of 1×10^5^, 5×10^5^, 1×10^6^, and 5×10^6^ cells/mL in PBS containing 0.1% BSA. For initial testing, 50 μL aliquots of 1×10^5^ cells/mL suspensions were dispensed into 384-well plates in triplicate using both regular chips and high recovery chips, with approximately 100 cells sorted per replicate. Higher concentration samples (5×10^5^-5×10^6^ cells/mL) were processed using regular chips only, with 384 cells sorted per replicate. Large-scale sorting (10,000 HeLa cells) was performed by dispensing 50 μL of 1×10^6^ cells/mL suspension into 384-well plates using regular chips, with PBS washing between sorting cycles. Approximately 26 plates underwent single-cell sorting for analysis. Lymph cells were similarly diluted to 1×10^5^-5×10^6^ cells/mL in PBS/0.1% BSA, while brain and intestine cells were prepared at 1×10^5^-1×10^6^ cells/mL in both HBSS/0.1% BSA and PBS. Lymph cells, brain cells and intestine cells were subjected to identical sorting protocols as HeLa cells. For recovery rate assessment, 5 μL aliquots of 1×10^5^ cells/mL tissue suspensions were processed through both chip types until complete dispensing.

### LC-MS/MS analysis

LC-MS/MS analysis was performed on either an Orbitrap Astral or Orbitrap Astral Zoom mass spectrometer (Thermo Fisher Scientific), each coupled to a Vanquish Neo UHPLC system. Peptide separation was carried out on an Aurora Elite TS analytical column (15 cm× 75 μm or 5 cm× 75 μm, IonOpticks), which was directly connected to an EASY-Spray (for 15 cm× 75 μm analytical column) or Flex-Spray (for 5 cm× 75 μm analytical column) ion source. The mobile phases consisted of solvent A (0.1% formic acid in water) and solvent B (80% acetonitrile with 0.1% formic acid).

For the 14-min gradient condition using the Orbitrap Astral Zoom mass spectrometer, peptide separation was performed on a 15 cm× 75 μm analytical column. Approximately11 min were dedicated to sample loading and post-run column equilibration, resulting in a total cycle time of ∼25 min. The LC gradient was configured as follows: at a flow rate of 0.45 μL/min, mobile phase B increased from 1% to 4% in 0.1 min and from 4% to 12% over the next 1.8 min. The flow rate was then reduced to 0.2 μL/min, holding B at 12% for 0.1 min, followed by a ramp to 28% over 6 min and to 40% over the next 3.5 min. B was then increased to 99% in 1 min and maintained for 1.5 min. To improve analytical performance and minimize background interference, a FAIMS Pro Duo interface was employed, with the compensation voltage set at −48 V. The Orbitrap MS1 scans were acquired at a resolution of 240,000 with a normalized AGC target of 500%, maximum injection time of 100 ms, RF lens set to 45%, and a scan range of 400-800 m/z. Data-independent acquisition (DIA) was performed using Astral MS2 scans, with an Auto DIA window scheme spanning a precursor mass range of 400-800 m/z and employing m/z Range mode for window placement. Fragmentation was achieved using higher-energy collisional dissociation (HCD) with a normalized collision energy of 25%. MS2 scans were collected over an m/z range of 150–2000, with the RF lens set to 45%, normalized AGC target of 800%, a DIA isolation window of 20 Th, a maximum injection time of 40 ms, and a loop control time of 0.6 seconds.

Under the 10.5-min gradient setting using the Orbitrap Astral mass spectrometer, peptide separation was carried out on the 15 cm× 75 μm analytical column and sample analysis was performed with approximately 9min dedicated to sample loading and post-run column equilibration, resulting in a total cycle time of 19min. The LC gradient was programmed as follows: at an initial flow rate of 0.75 μL/min, solvent B was increased from 6% to 12% within 0.9min. The flow rate was then reduced to 0.2 μL/min, with a gradual increase from 12% to 35% B over 7 min, followed by a rise to 50% B over 1 min. Subsequently, the flow rate was restored to 0.75 μL/min to ramp from 50% to 99% B in 0.5 min, with an isocratic hold at 99% B for an additional 1 min at a flow rate of 1.2 μL/min. MS/MS settings were the same as those described in 14-min gradient setting, except that the Orbitrap MS1 scan range was set to 400-900 m/z.

For the 5-min LC gradient setup by the Orbitrap Astral mass spectrometer, using a 15 cm× 75 μm analytical column and approximately 5 min allocated for sample loading and post-run column equilibration, resulting in a total cycle time of ∼9.5 min. The chromatographic program initiated at a flow rate of 0.5 μL/min, increasing solvent B from 2% to 12% within 0.5 min. The flow rate was then reduced to 0.25 μL/min to achieve a stepwise gradient from 12% to 12.2% B in 0.05 min, followed by a linear increase to 25% over 2.2 min and to 50% over the next 1.8 min. The gradient then transitioned rapidly to 99% B in 0.2 min at 0.75 μL/min, followed by an isocratic hold at 99% B for 0.25 min at 1.2 μL/min. MS1 and DIA settings were the same as those described in 14-min gradient setting, except that the FAIMS Pro Duo interface was employed, with the compensation voltage set at −52 V and the precursor scan range was adjusted to 400–840 m/z.

To enable high-throughput analysis in dual-spray TDI system, the Orbitrap Astral mass spectrometer was integrated with the Vanquish Neo UHPLC system, configured with two independent analytical pumps and a single autosampler. This setup allows the parallel operation of two identical separation columns, coordinated by an intelligent workflow and supported by a custom-built column controller capable of automated column switching. Two 5 cm× 75 μm analytical columns were employed for dual-spray mode. Two methods were implemented in dual-spray TDI system: a longer gradient for bulk proteomics and a shorter gradient optimized for SCP. The longer method had an effective separation time of 10.6 min. The column-switching process, including delay and alignment time, required approximately 1 min, resulting in a total cycle time of 12.2 min per run. The LC gradient was delivered at a flow rate of 0.25 μL/min, with mobile phase B increasing linearly from 4% to 40%. The shorter method had an effective separation time of 7.6 min, with a total cycle time of 8.6 min. The LC gradient was delivered at a flow rate of 0.4 μL/min, with mobile phase B increasing linearly from 1% to 35%. MS/MS settings were the same as those described in 14-min gradient setting, except that the compensation voltage of FAIMS Pro Duo interface was set to −49 V.

### Data analysis

To evaluate LC-MS system stability, quality control (QC) samples comprising 250 pg of HeLa cell digest (Pierce™) and column blanks (0.1 μL of 0.1% formic acid in LC-MS–grade water, v/v) were applied. Raw spectral data were processed with DIA-NN (v1.9.2) in library-free mode against the human UniProt reference proteome (2024 release) supplemented with 246 common contaminant sequences. Mass and MS1 accuracy were set to 0 (auto mode). “Heuristic protein inference” and “match-between-runs (MBR)” were enabled. The maximum number of variable modifications was limited to two. Carbamidomethylation of cysteine was not included.

For SCP, raw files were analyzed using a library-free directDIA workflow in Spectronaut v19 (Biognosys) against the mouse UniProt reference proteome (2024 release) plus the same contaminant set. Variable modifications included methionine oxidation and N-terminal acetylation; carbamidomethylation of cysteine was excluded. Cross-run normalization was disabled.

Protein-by-sample quantification tables (.tsv files) exported from Spectronaut were imported into R and used to construct a Seurat object. All downstream analyses were performed using the Seurat package (v5)^45^. Cells with more than 100 identified proteins were retained for further analysis.

Tissue specific marker proteins were selected using the FindAllMarkers function. Heatmap visualization of tissue specific marker proteins were performed using the ComplexHeatmap package, based on log normalized protein intensities and the rows were scaled^46^. Gene Ontology (GO) enrichment analysis was conducted within tissue specific marker proteins, focusing on the Biological Process (BP) category. The analysis was performed using the clusterProfiler package, and significantly enriched GO terms (adjusted p < 0.05) were identified^47^.

Among the tissues analyzed, only the kidney sample was acquired on different days. To account for potential batch effects arising from technical variability, kidney cells from each loading batch were initially processed independently and subsequently integrated. Normalization was performed using NormalizeData, and highly variable proteins were selected using the variance-stabilizing transformation (VST) method via the FindVariableFeatures function. Each batch-specific dataset was scaled with ScaleData and subjected to PCA using the top 30 principal components (RunPCA). To integrate the datasets, integration anchors were identified using FindIntegrationAnchors with reduction = “rpca”, followed by data integration using the IntegrateData function. This yielded a unified expression matrix stored in a new assay labeled “integrated”. The samples from the remaining tissues were processed according to the process of normalization, selecting highly variable proteins, scaling, PCA analysis, unsupervised clustering, and t-SNE dimensionality reduction. Cellular subtype annotation was based on significantly enriched genes in each cluster using the FindAllMarkers function. To further validate, the expression of canonical marker proteins was showed in violin plots.

Pseudotime analysis was performed using the Slingshot package^48^. Cells were ordered along inferred trajectories based on annotated clustering and dimensionality-reduced coordinates. Marker protein expression was plotted along pseudotime to reveal stage-specific dynamics.

### Data availability

The MS proteomics data have been deposited to the ProteomeXchange Consortium (http://proteomecentral.proteomexchange.org) via the iProX partner repository with the dataset identifier PXD064964.

## Supporting information

End-to-end high-throughput single-cell proteomics via SPRINT and dual-spray LC-MS

## Acknowledgements

This work was supported by funds from the Non-profit Central Research Institute Fund of the Chinese Academy of Medical Sciences (grant no. 2023-RC180-03), the Chinese Academy of Medical Sciences (CAMS) Innovation Fund for Medical Sciences (grant no. 2022-I2M-2-004, 2023-I2M-2-005, 2021-I2M-1-061), National Natural Science Foundation of China (22574120), Natural Science Foundation of Jiangsu Province (BK20240443), the Suzhou Municipal Key Laboratory (SZS2022005) and the NCTIB Fund for the R&D Platform for Cell and Gene Therapy.

## Author Contributions

Z.Y. conceptualized the study. Z.L., W.D and L.G. performed most of the experiments and data analysis. R.G., J.D., H.Z. contributed to optimizing and performing the single-cell analysis. X.Z. performed part of data analysis. All authors read, edited and approved the final version of the manuscript.

