## Supplementary material for "End-to-end high-throughput single-cell proteomics via SPRINT and dual-spray LC-MS": End-to-end high-throughput single-cell proteomics via SPRINT and dual-spray LC-MS

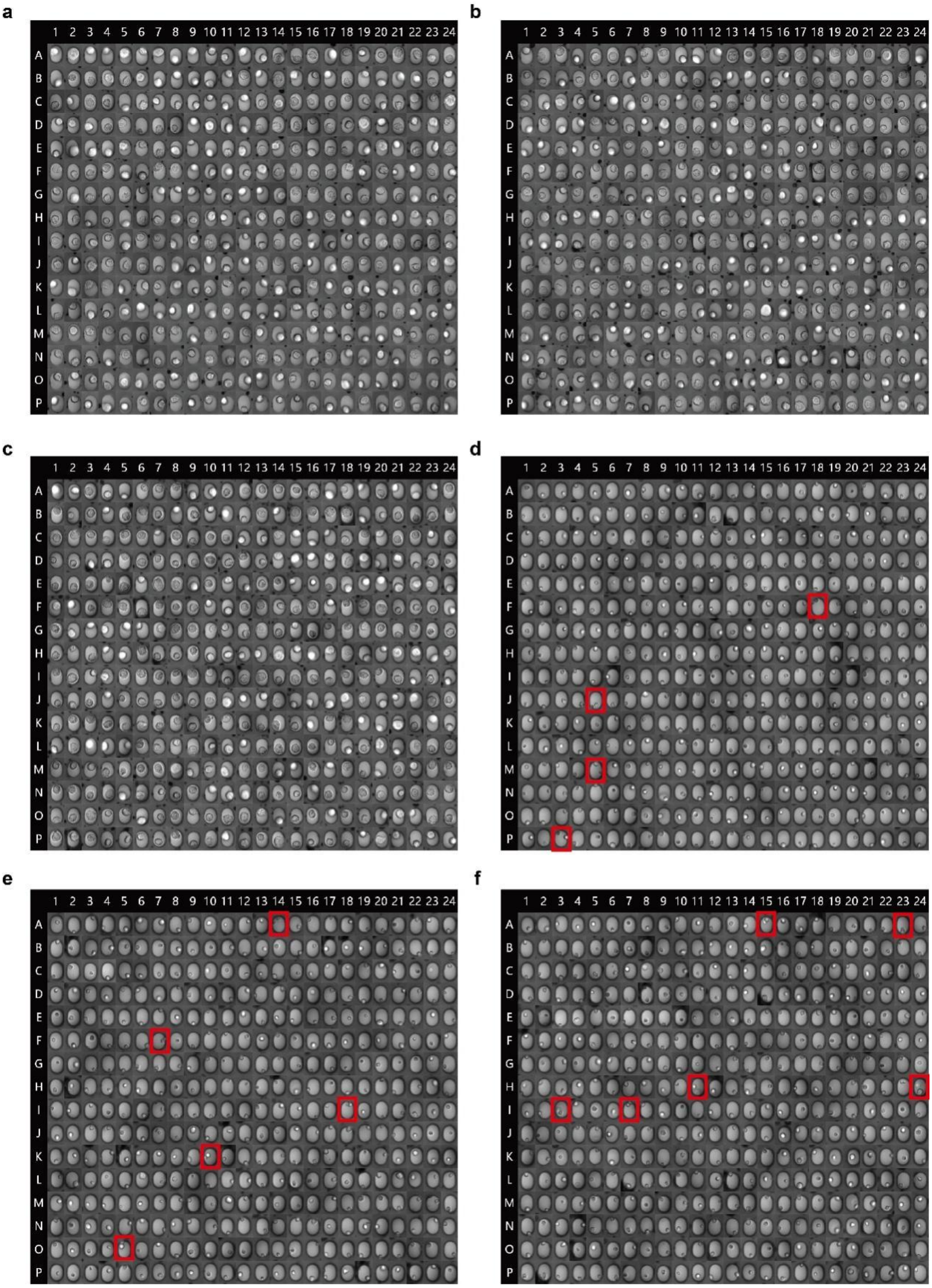


**Figure S1 | Single-cell sorting accuracy for HeLa and bone marrow cells.** HeLa cells (**a-c**) and bone marrow cells (**d-f**) were sorted into 384-well plates in triplicate using regular chips. Post-sorting, the wells were manually inspected to evaluate the presence of debris, single cells, and doublets. Wells containing debris or doublets are marked with red boxes. Sorting accuracy was calculated as the ratio of wells containing a single cell to the total number of wells (384).

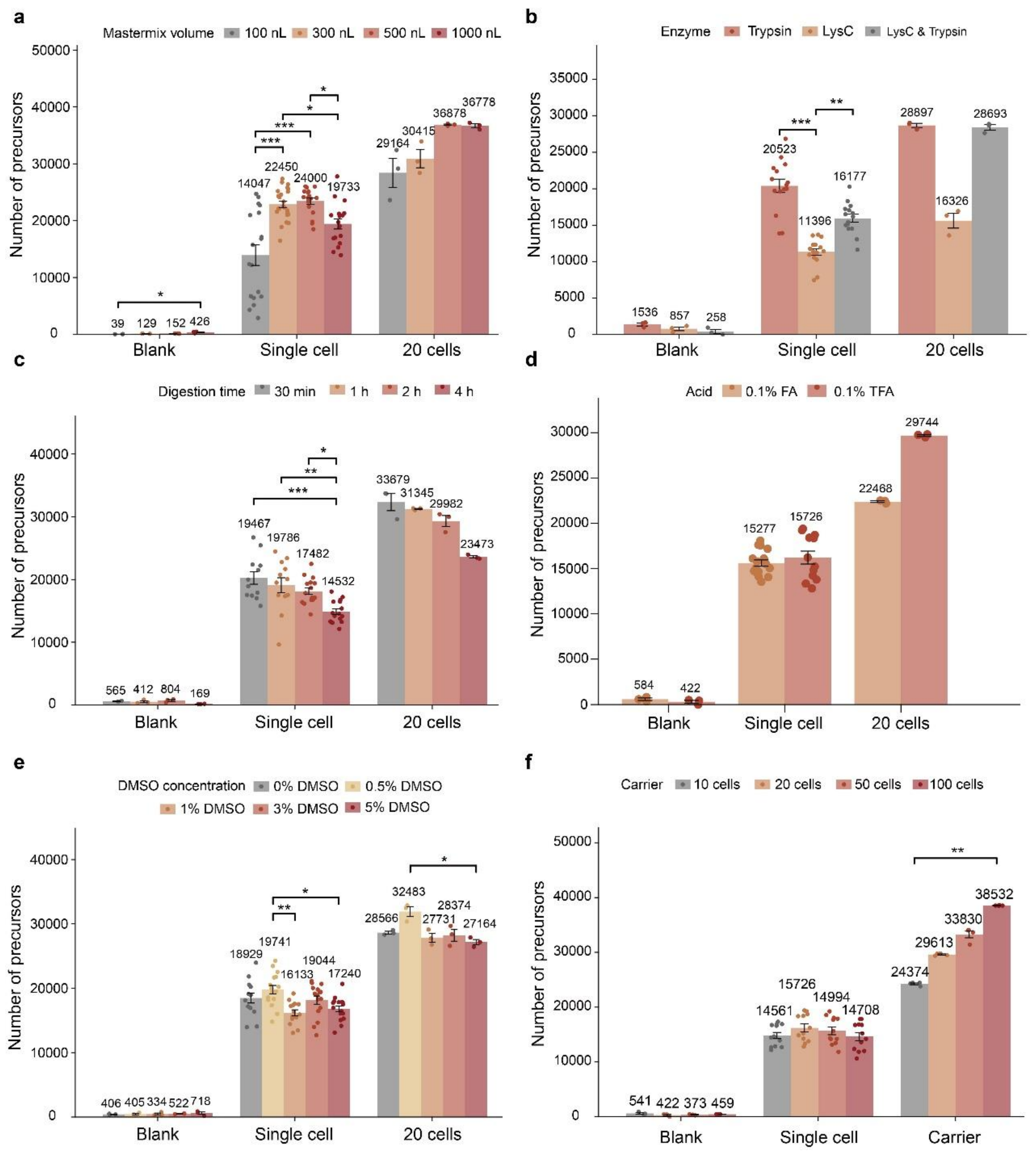


**Figure S2 | Enhanced precursor identifications depth through integrated workflow optimization.** (**a**) Precursor identifications across different master mix volumes. (**b**) Effect of enzyme type on precursor identifications. (**c**) Impact of digestion time on precursor identifications. (**d**) Comparison of 0.1% FA versus 0.1% TFA in 1% DMSO buffer. (**e**) Effect of DMSO concentration on precursor identifications. (**f**) Effect of carrier cell number on precursor identifications. Data represents HeLa single-cell or 20-cells analyses, with blank as controls. *, $p < 0.05$; **, $p < 0.01$; ***, $p < 0.001$. Except for panel d, which employed the Wilcoxon rank-sum test for pairwise comparisons, all other panels (a-c, e-f) used Dunn's test following Kruskal-Wallis analysis. *p*-values were adjusted using the Bonferroni method.

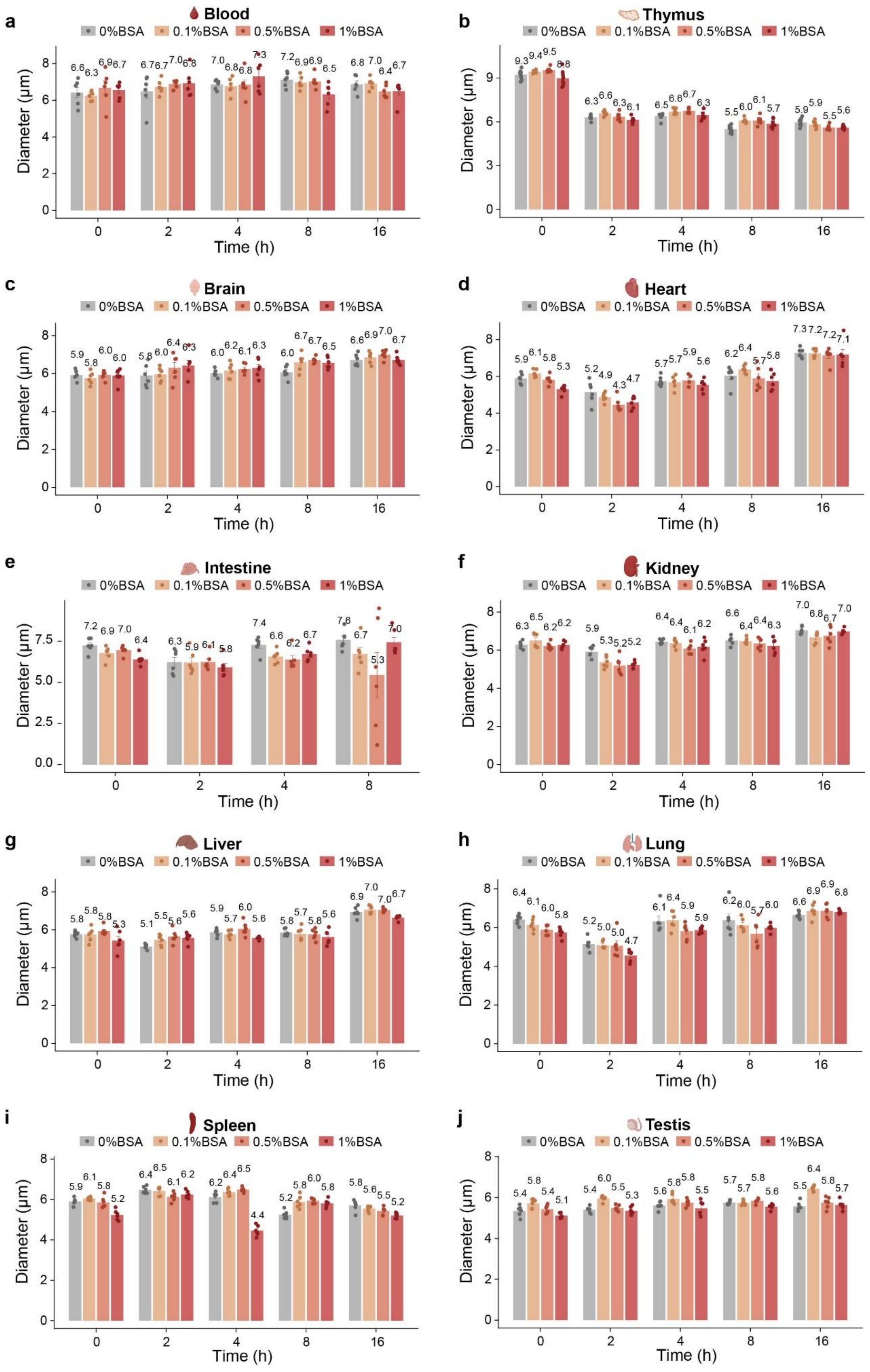


**Figure S3 | Cell diameter changes over time for different tissue types across tested conditions.** (**a-j**) Cell diameter dynamics of ten tissue types: blood (**a**), thymus (**b**), brain (**c**), heart (**d**), intestine (**e**), kidney (**f**), liver (**g**), lung (**h**), spleen (**i**) and testis (**j**). Cell diameter was measured at 0, 2, 4, 8 and 16 h under different BSA concentrations using an automated cell counter.

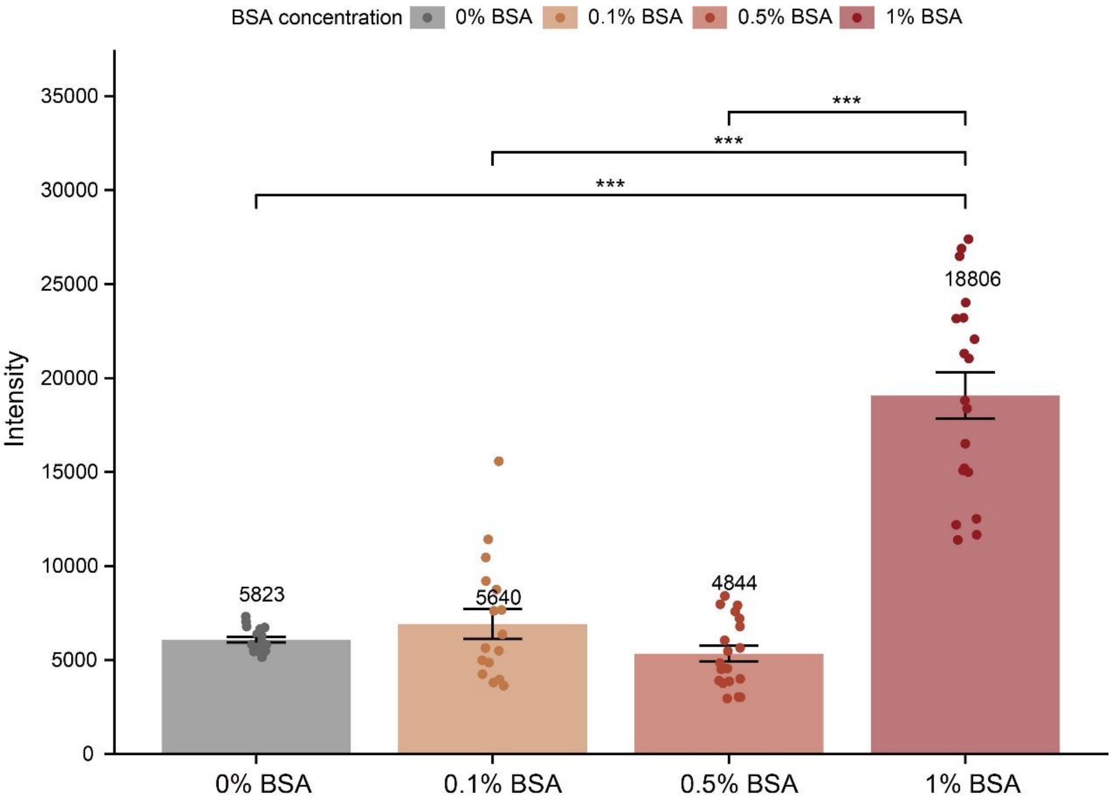


**Figure S4 | The BSA intensity of single-cell proteomics under different BSA concentrations.** HeLa single cells suspended in PBS containing varying BSA concentrations (0%, 0.1%, 0.5% and 1%) were analyzed using the Vanquish Neo–Orbitrap Astral system. The raw data were processed with a library-free directDIA workflow in Spectronaut v19, and BSA intensity values were obtained from the protein-by-sample quantification tables (.tsv files). *, $p < 0.05$; **, $p < 0.01$; ***, $p < 0.001$. The panel used Dunn's test following Kruskal-Wallis analysis. *p*-values were adjusted using the Bonferroni method.

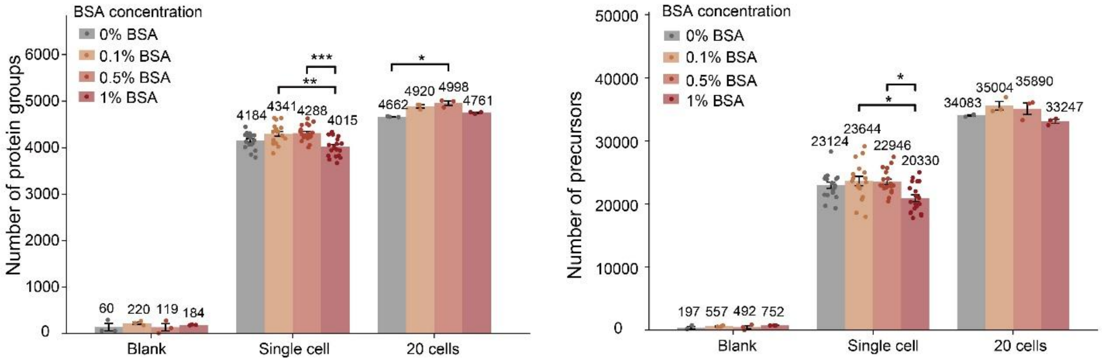


**Figure S5 | Protein and precursor identifications under different BSA concentrations.** HeLa cells suspended in PBS with varying BSA concentrations (0%, 0.1%, 0.5% and 1%) were analyzed on the Vanquish Neo–Orbitrap Astral system. The raw data were processed using a library-free directDIA workflow in Spectronaut v19, and protein and precursor identifications was obtained from the output reports. Data represents HeLa single-cell or 20-cell analyses, with blank as controls. *, $p < 0.05$; **, $p < 0.01$; ***, $p < 0.001$. The panel used Dunn's test following Kruskal-Wallis analysis. $p$-values were adjusted using the Bonferroni method.

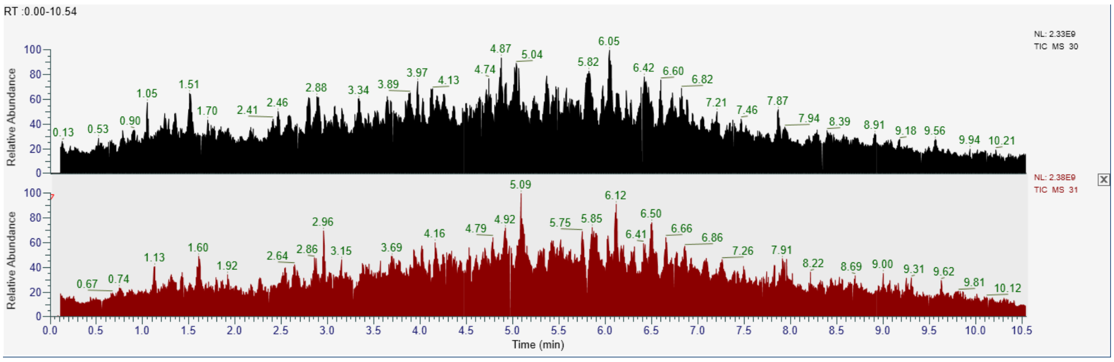


**Figure S6 | Representative total ion chromatogram (TIC) of two analytical columns in the dual-spray TDI system.** A total of 100 ng of HeLa peptides were analyzed using the dual-spray TDI system with a 12-min cycle method. The upper TIC corresponds to one analytical column, while the lower TIC corresponds to the other analytical column.

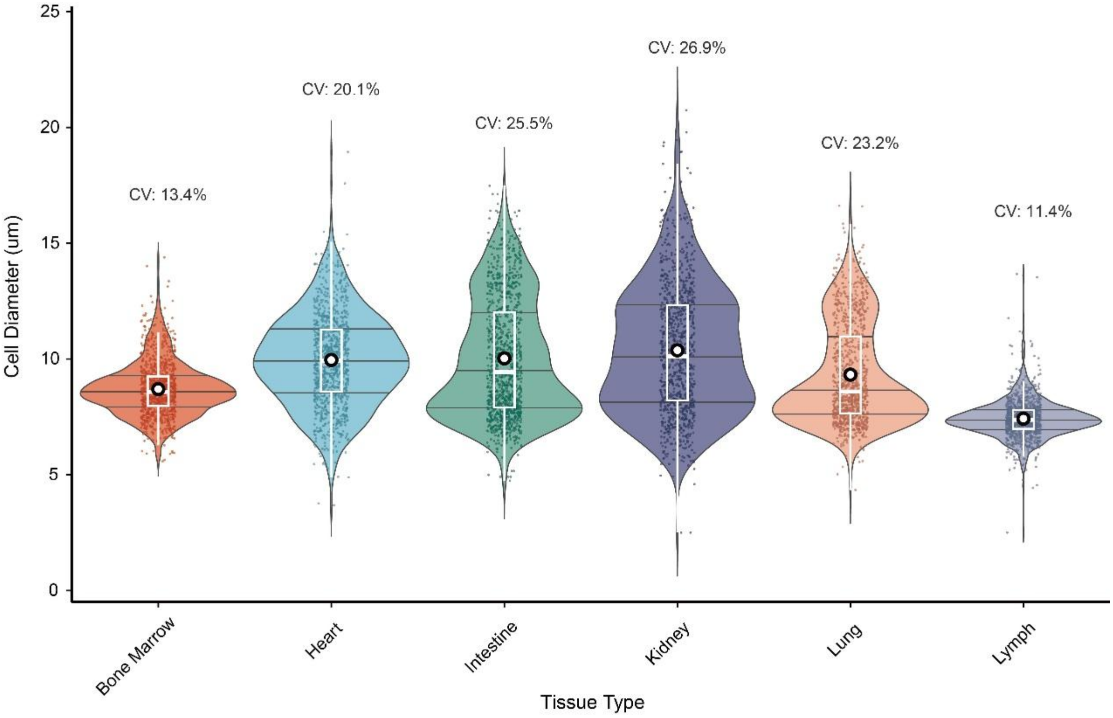


**Figure S7 | Violin plot of cell diameter distributions across six tissue types.** Single cells from bone marrow, heart, intestine, kidney, lung and lymph tissue were sorted using the SPRINT platform. Cell diameter measurements were obtained from the system output reports.

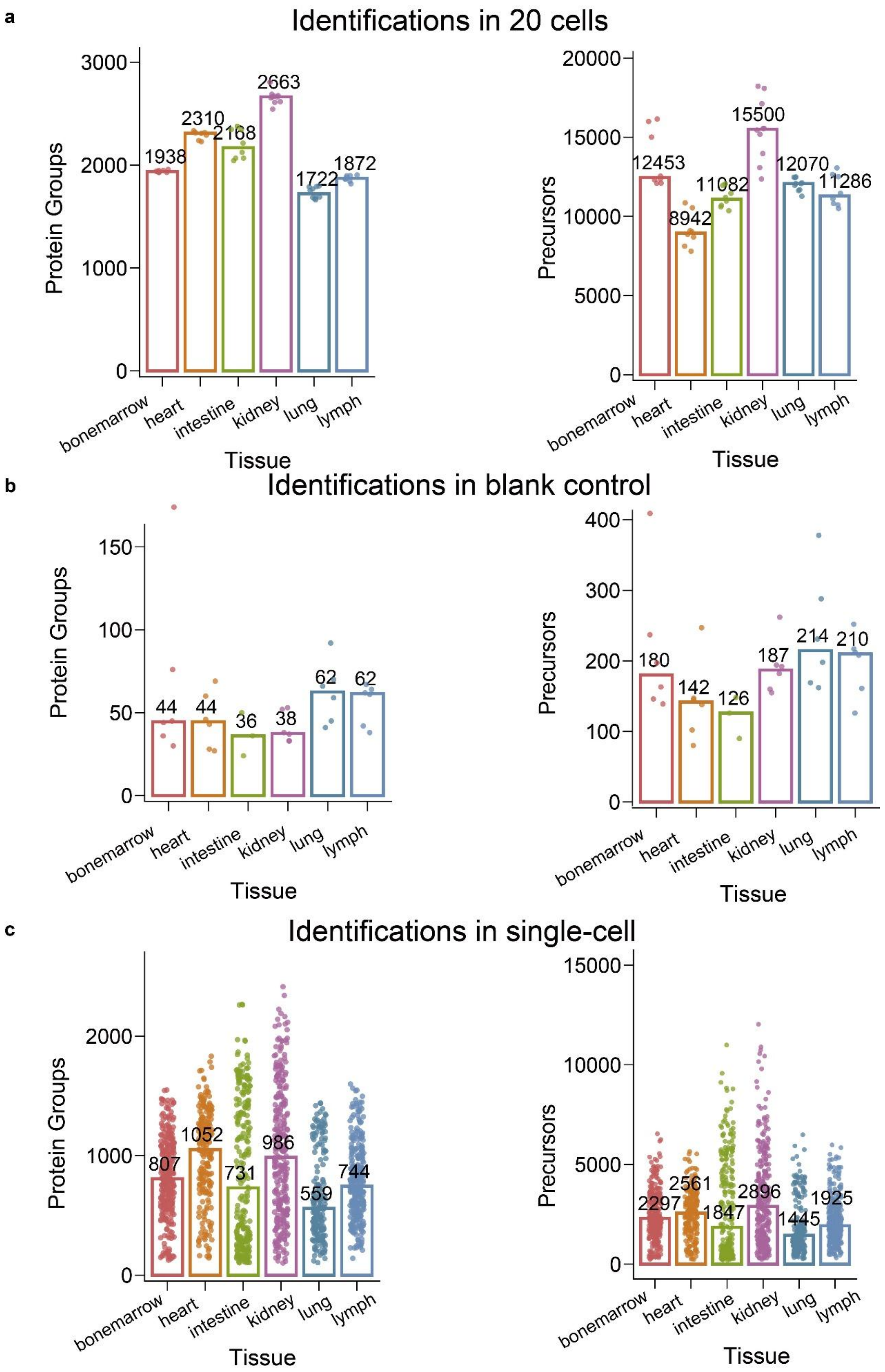


**Figure S8 | The protein group and precursor identifications across six tissue types.** Single cells from bone marrow, heart, intestine, kidney, lung and lymph tissue were sorted using the

SPRINT platform and analyzed with the dual-spray TDI system. Protein group and precursor identifications are shown for 20 pooled cells (**a**), blank controls (**b**), and single-cells (**c**).

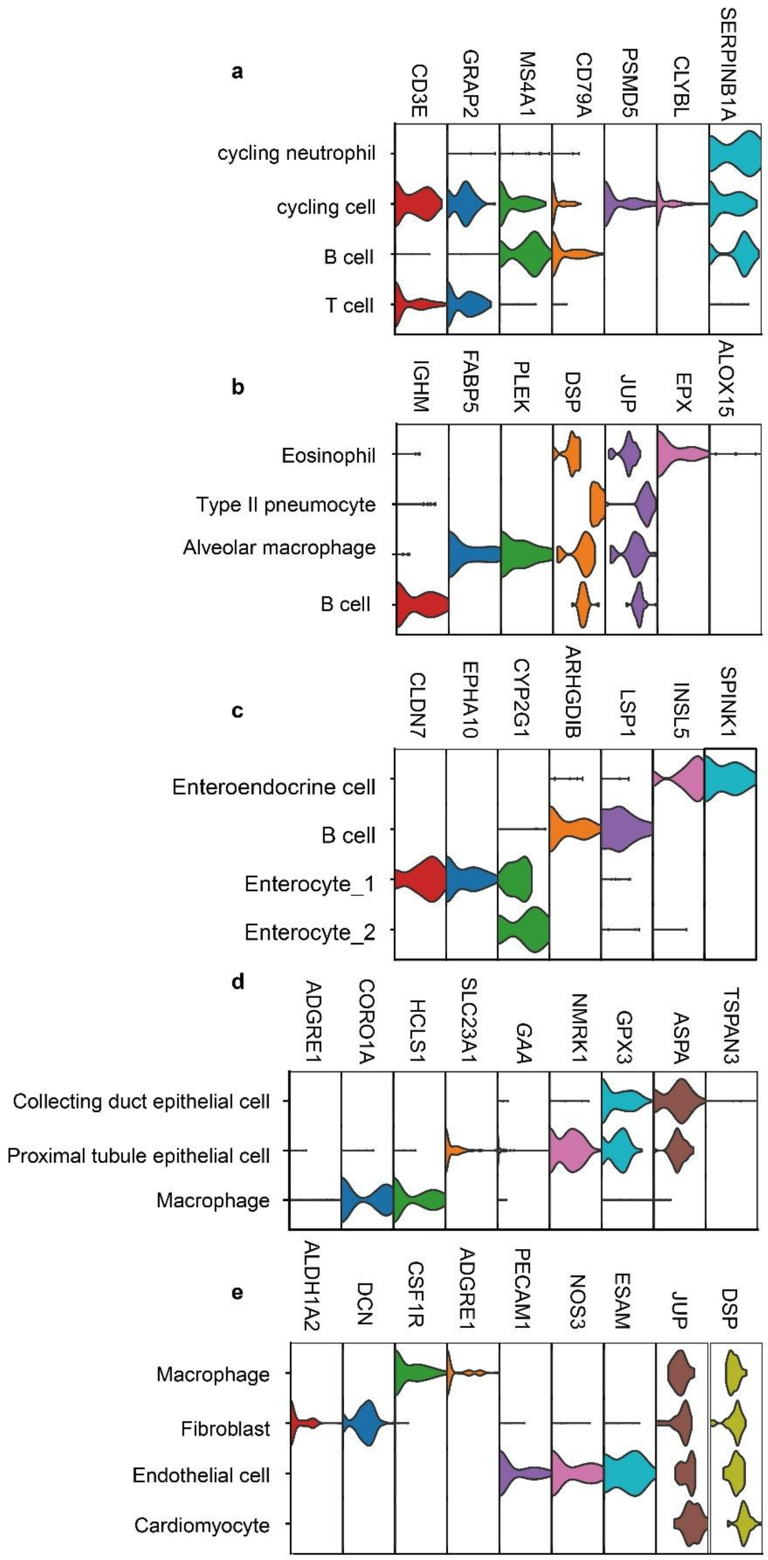


**Figure S9 | Violin plots showing canonical marker protein expression across cell subclusters.** (**a-e**) Cell subclusters from five tissue types: lymph (**a**), lung (**b**), intestine (**c**), kidney (**d**), and heart (**e**). Violin plots were generated based on the expression levels of canonical marker proteins within each subcluster.

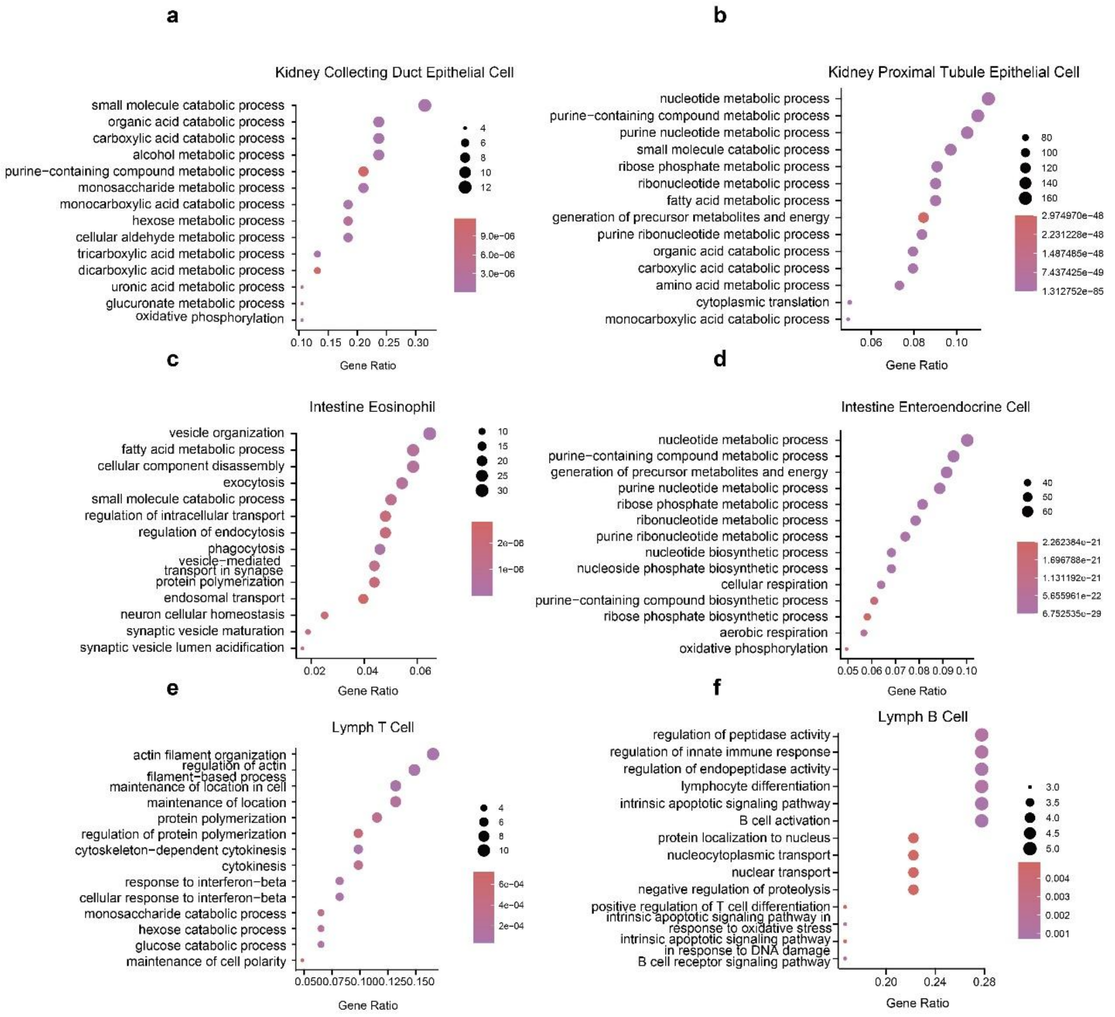


**Figure S10 | The GO analysis of cell subclusters deprived from kidney, intestine and lymph tissues.** (**a-f**) Cell subclusters from three tissue types: kidney collecting duct epithelial cell (**a**), kidney proximal tubule epithelial cell (**b**), intestine eosinophil (**c**), intestine enteroendocrine cell (**d**), lymph T cell (**e**), and lymph B cell (**f**). GO enrichment analysis was performed on tissue-specific marker proteins, focusing on the Biological Process (BP) category. Significantly enriched GO terms (adjusted $p < 0.05$) were identified using the clusterProfiler package.

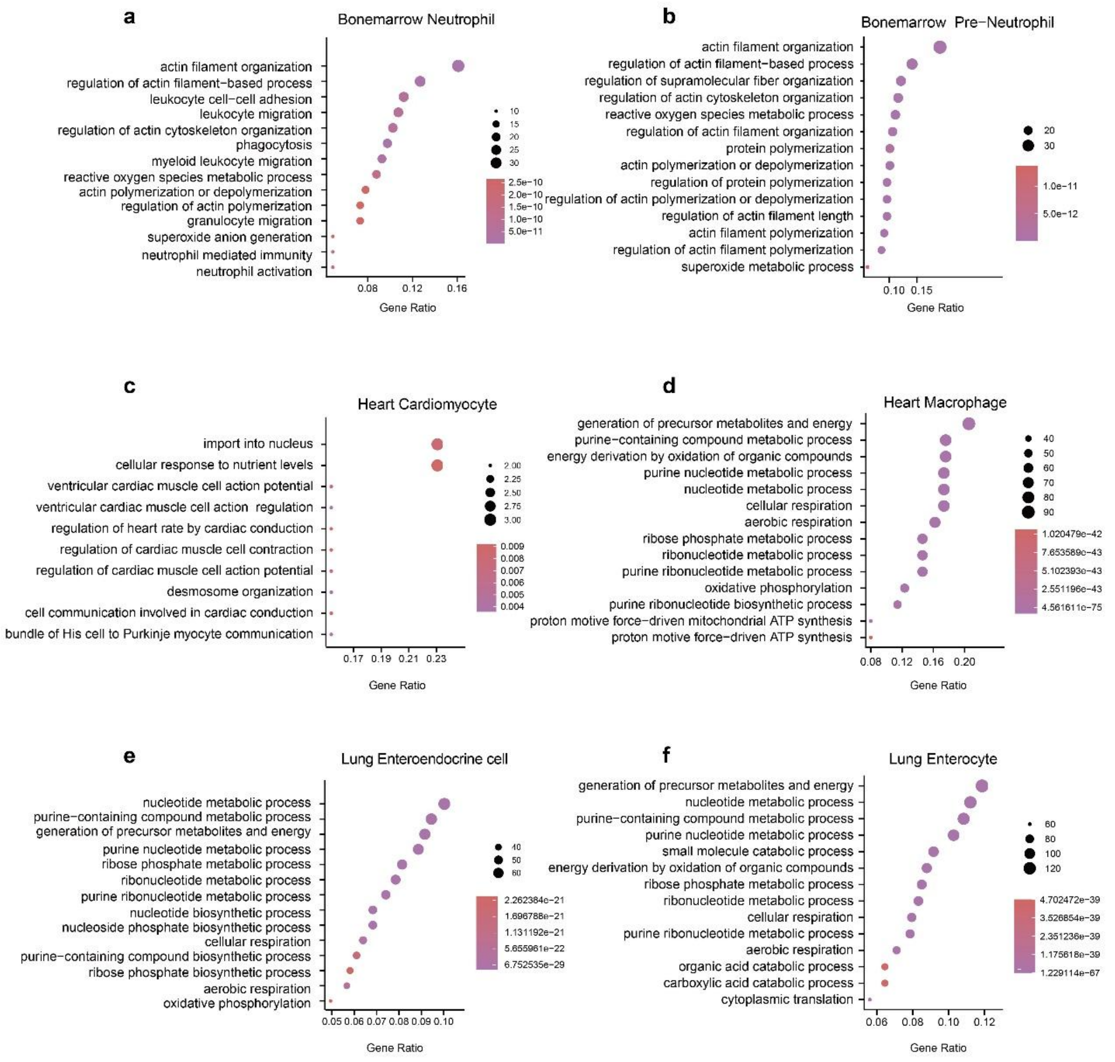


**Figure S11 | The GO analysis of cell subclusters deprived from bone marrow, heart and lung tissues.** (**a-f**) Cell subclusters from three tissue types: bone marrow neutrophil (**a**), bone marrow pre-neutrophil (**b**), heart cardiomyocyte (**c**), heart macrophage (**d**), lung enteroendocrine cell (**e**), and lung enterocyte (**f**). GO enrichment analysis was performed on tissue-specific marker proteins, focusing on the Biological Process (BP) category. Significantly enriched GO terms (adjusted $p < 0.05$) were identified using the clusterProfiler package.
